# Targeting the Hippo pathway in cancers via ubiquitination dependent TEAD degradation

**DOI:** 10.1101/2023.10.03.560675

**Authors:** Trang H. Pham, Kanika Bajaj Pahuja, Thijs J. Hagenbeek, Jason Zbieg, Cameron L. Noland, Victoria C. Pham, Xiaosai Yao, Christopher M. Rose, Kristen C. Browder, Ho-June Lee, Mamie Yu, May Liang-Chu, Scott Martin, Erik Verschueren, Jason Li, Marta H. Kubala, Rina Fong, Maria Lorenzo, Paul Beroza, Peter Hsu, Sayantanee Paul, Elisia Villemure, Wendy Lee, Tommy K. Cheung, Saundra Clausen, Jennifer Lacap, Yuxin Liang, Jason Cheng, Steve Schmidt, Zora Modrusan, Michael Cohen, James Crawford, Heinrich Jasper, Alan Ashworth, Jennie R. Lill, Shiva Malek, Joachim Rudolph, Ingrid E. Wertz, Matthew T. Chang, Xin Ye, Anwesha Dey

## Abstract

The Hippo pathway is among the most frequently altered key signaling pathways in cancer. TEAD1-4 are essential transcription factors and key downstream effectors in the Hippo pathway in human cells. Here, we identified RNF146 as a ubiquitin ligase (E3) of TEADs, which negatively regulates their stability in cells through proteasome-mediated degradation. We show that RNF146-mediated TEAD ubiquitination is dependent on the TEAD PARylation state. We further validated the genetic interaction between RNF146 and the Hippo pathway in cancer cell lines and the model organism *Drosophila melanogaster.* Despite the RNF146 and proteasome-mediated degradation mechanisms, TEADs are stable proteins with a long half-life in cells. We demonstrate that degradation of TEADs can be greatly enhanced pharmacologically with heterobifunctional chemical inducers of protein degradation (CIDEs). These TEAD-CIDEs can effectively suppress activation of YAP/TAZ target genes in a dose-dependent manner and exhibit significant anti-proliferative effects in YAP/TAZ-dependent tumor cells, thus phenocopying the effect of genetic ablation of TEAD protein. Collectively, this study demonstrates that the ubiquitin-proteasome system plays an important role in regulating TEAD functions and provides a proof-of-concept demonstration that pharmacologically induced TEAD ubiquitination could be leveraged to target YAP/TAZ-driven cancers.

## INTRODUCTION

The genetic alterations of Hippo pathway components components have been observed in 10% of human cancers, including glioma, breast, liver, lung, prostate, colorectal, and gastric cancers (Dong et al., 2007; Lo Sardo et al., 2018; Sanchez-Vega et al., 2018; Steinhardt et al., 2008; Zhao et al., 2007). In addition, non-genetic dysregulations of the pathway are broadly involved in tumor development (Calses et al., 2019; Dey et al., 2020; Wang et al., 2018; Zanconato et al., 2016). These upstream dysfunctions converge on TEAD transcription factors as a central node to regulate transcriptional output driving tumor progression, metastasis, immunity, cancer metabolism, and drug resistance (Cordenonsi et al., 2011; Halder and Johnson, 2011; Kapoor et al., 2014; Lin et al., 2015; Yu and Guan, 2013). There are four TEAD members in humans, TEAD1 (TEF-1/NTEF), TEAD2 (TEF-4/ETF), TEAD3 (TEF-5/ETFR-1), and TEAD4 (TEF-3/ETFR-2/FR-19) (Lin et al., 2017; Yasunami et al., 1996) that share high structural similarity and a conserved palmitoylation pocket (Calses et al., 2019; Huh et al., 2019; Lin et al., 2017). Even though TEADs are mostly chromatin-bound in cells (Li et al., 2015), they do not exhibit any transcriptional activity on their own and rely on their coactivators to turn on gene expression (Xiao et al., 1991). Conversely, the two main Hippo pathway transcription effectors, YAP (*YAP1*) and TAZ (*WWTR1*), control gene expression via interacting with TEADs at their C-terminal transactivating domains (Vassilev et al., 2001; Zhang et al., 2009; Zhao et al., 2008).

Consistent with TEADs’ central roles in controlling the Hippo pathway output, genetic studies revealed that TEADs play important roles during development, including cardiogenesis, neural development, and trophectoderm lineage determination (Chen et al., 1994; Huh et al., 2019; Kaneko et al., 2007; Nishioka et al., 2009; Nishioka et al., 2008; Yagi et al., 2007; Yasunami et al., 1996). Furthermore, numerous studies underline the importance of TEADs in human cancers. Overexpression of TEADs has been implicated in multiple stages of cancer progression across cancer types. Moreover, the Hippo pathway has been shown as a common mechanism of resistance to chemo- and targeted therapies (Kapoor et al., 2014; Lin et al., 2015; Song et al., 2015). To date, multiple efforts to target the Hippo pathway in cancers have mainly focused on small molecules binding to the palmitoylation pocket of TEAD, which can subsequently block the interaction between YAP/TAZ and TEADs (Calses et al., 2019; Dey et al., 2020).

Recent innovations in the field of targeted protein degradation and manipulation of the ubiquitin-proteasome system have enabled new therapeutic approaches (Scheepstra et al., 2019). Targeted protein degradation using heterobifunctional chemical inducers of protein degradation (CIDEs), also known as PROTACs (Sakamoto et al., 2001) has emerged as a new frontier of drug discovery (Chamberlain and Hamann, 2019; Yang et al., 2019). Such heterobifunctional degraders consist of two moieties, one binding to the target protein and the other to an E3 ubiquitin ligase to induce proximity between the E3 ubiquitin ligase with the target, thus causing ubiquitination and subsequent proteasomal degradation of the target protein. Such degrader molecules offer several advantages over small molecule inhibitors. They act through an event-driven mechanism instead of conventional occupancy-driven pharmacology; in addition, target protein degradation removes all functions of the target protein and may also lead to the destabilization of entire multidomain protein complexes (Chamberlain and Hamann, 2019; Yang et al., 2019). Heterobifunctional degraders have been reported for a diverse array of oncoproteins, and some have entered clinical trials (Han et al., 2019; You et al., 2020).

In this study, we explored intrinsic and induced TEAD degradation via the ubiquitin-proteasome system to suppress the function of TEAD proteins. We identified PARylation-dependent ubiquitin ligase RNF146 as an endogenous ubiquitin ligase for TEAD transcription factors. We further demonstrate that RNF146 is a potent negative regulator of TEAD transcriptional activity, indicating that TEAD ubiquitination can be explored to suppress the Hippo target gene activation upon YAP/TAZ stabilization (Fan et al., 2020). To this end, we report the discovery and characterization of a TEAD-CIDE, Compound D, that can induce potent and sustained degradation of all four human TEAD isoforms by co-opting the CRL4-CRBN ubiquitin ligase complex. Compound D exhibits significant inhibition of cell proliferation and downstream signaling compared to a reversible TEAD lipid pocket binder that lacks the ubiquitin ligase binding moiety. As a proof of concept, we show that TEAD-CIDE-induced degradation had anti-proliferation effects comparable to genetic TEAD depletion mediated by sgRNA or siRNA in Hippo pathway-dysregulated cell lines. Collectively, our study demonstrates that endogenous ubiquitination of TEADs via RNF146 or induced ubiquitination by TEAD-CIDE is an effective mechanism to suppress the activation of the Hippo pathway target genes. This work highlights previously unrecognized TEAD ubiquitination and PARylation crosstalk and a new strategy for targeting the Hippo pathway in cancers.

## RESULTS

### TEAD transcription factors are ubiquitinated in cells

To assess post-translational modifications that could impact TEAD stability, particularly ubiquitination, we transfected HEK-293 cells with expression vectors for each of the four TEAD paralogs, followed by a cell-based ubiquitination assay. Our results indicated that all four paralogs can be ubiquitinated in cells (Figure 1A), with TEAD2 and TEAD4 showing the highest levels of poly-ubiquitination. Focusing on TEAD4, we found that its ubiquitination levels were further increased upon treatment with proteasome inhibitor MG132 in a time-dependent manner (Figure 1B), suggesting that TEAD4 ubiquitination can result in proteasomal degradation. To further assess the ubiquitination of TEADs, we performed a peptide level immunoaffinity enrichment (PTMScan) experiment to evaluate the profiles of K-GG peptides (the result of trypsin digestion of ubiquitinated proteins) for ectopically expressed TEAD4 in HEK293 cells (Figure 1C). We identified several ubiquitination sites (Figure 1D) and mapped them on TEAD4 using known crystal structures. These ubiquitination sites did not reside in any specific TEAD structural domain (Figure S1A, B) (Mesrouze et al., 2017; Shi et al., 2017). Consistent with our cell-based ubiquitination assay (Figure 1B), the ubiquitinated peptides were significantly increased following MG132 treatment (Figure 1E, S1C and Supplemental Table 1). Taken together, our data demonstrates that the TEAD transcription factors can be poly-ubiquitinated in cells, and such poly-ubiquitination can target TEADs for proteasomal degradation.

**Figure 1.**
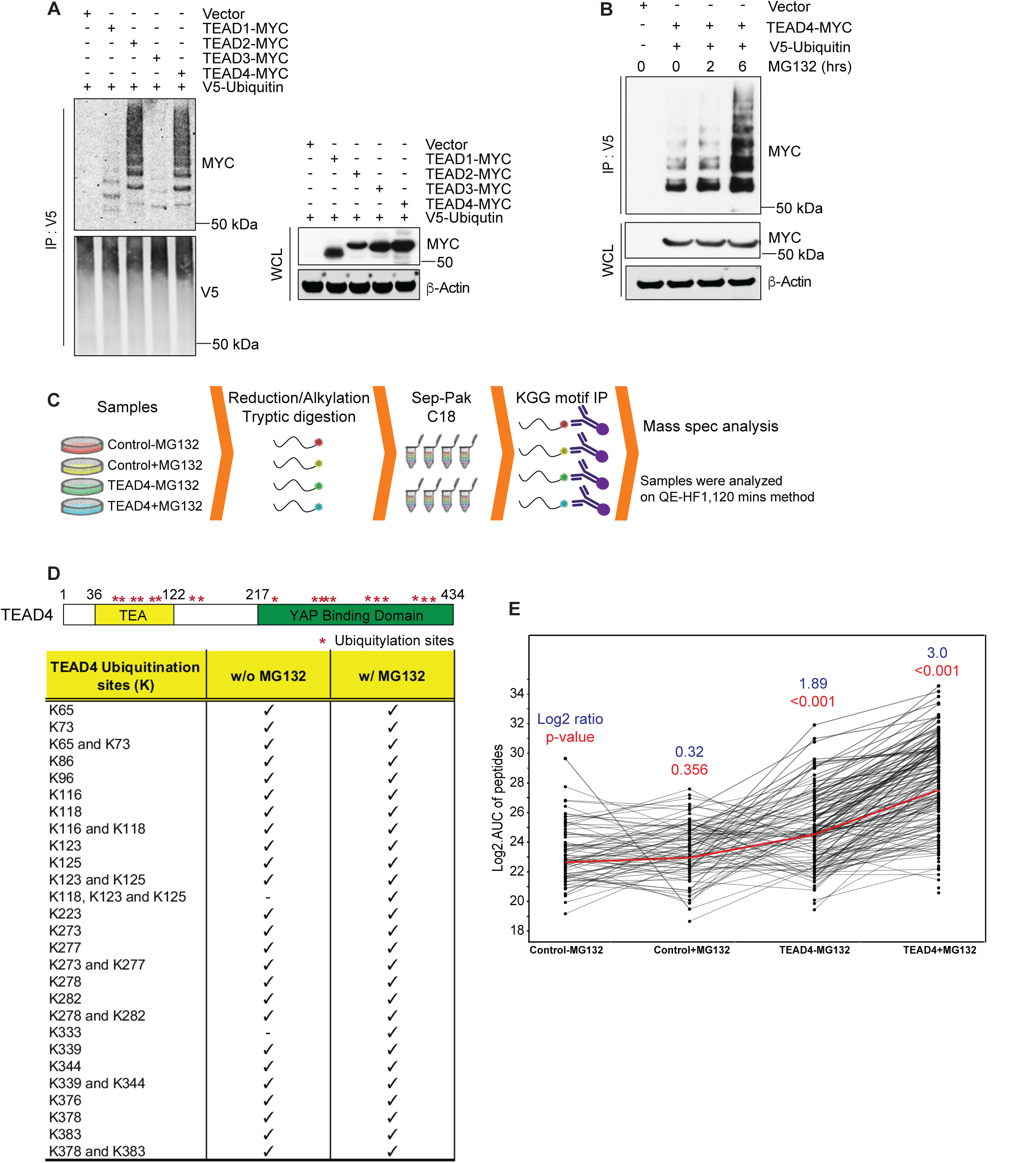
TEAD transcription factors are ubiquitinated and targeted for proteasome-dependent degradation. **(A)** TEADs are ubiquitinated. HEK293 cells were transfected with a control vector, TEAD1-Myc, TEAD2-Myc, TEAD3-Myc, or TEAD4-Myc and V5-ubiquitin plasmids for 48 hours, followed by immunoprecipitation with an anti-V5 antibody. TEAD expression and ubiquitinated TEAD were detected with anti-Myc antibody (n=3). **(B)** MG132 induces accumulation in TEAD ubiquitination. HEK293 cells were transfected with TEAD4-Myc and V5-ubiquitin plasmids for 48 hours, followed by immunoprecipitation with anti-V5 antibody after an additional treatment with 10 µM MG132. TEAD4 expression and ubiquitinated TEAD4 were detected with anti-Myc antibody (n=3). **(C-E)** Ubiquitination sites on TEAD4 were mapped using KGG motif immunoprecipitation, followed by LC-MS/MS. C, experimental schematic: TEAD4-Myc was overexpressed in HEK293 cells for 48 hours, MG132 was added 6 hours prior to cell lysis where indicated. KGG motif immunoprecipitation was then performed, followed by LC-MS/MS. D, each ubiquitination site was represented by * and summarized in the table. E, ubiquitination for TEAD4 across conditions. Red line represents the aggregate of individually quantified K-GG peptides (black lines) using a linear mixed-effect model. Each quantification event represents the non-redundant, absolute peak area (converted to log2) under a given condition for a confidently assigned K-GG peptide spectral match.

#### RNF146 is the E3 ligase for TEAD transcription factors and negatively regulates TEAD transcriptional activity

We next sought to identify the ligase responsible for TEAD ubiquitination. We performed a screen using a siRNA library for 640 ubiquitin-related genes in MCF7 cells that carry a luciferase reporter driven by multimerized TEAD binding sites (8xGTIIC-Luc). We identified three ubiquitin ligases – RNF146 and TRAF3 – as potential regulators of the Hippo pathway from the primary screen using the luciferase reporter (Supplemental Table 2 and Figure S2A). Next, we validated these hits by assessing their impact on the TEAD target genes CTGF and CYR61/CCNE1 upon siRNA-mediated gene knockdown (Figure S2B). Two independent siRNAs targeting RNF146 consistently enhanced the expression of both genes, again indicating *RNF146* as a putative negative regulator of TEAD-mediated gene expression (Figure 2A and S2B).

**Figure 2.**
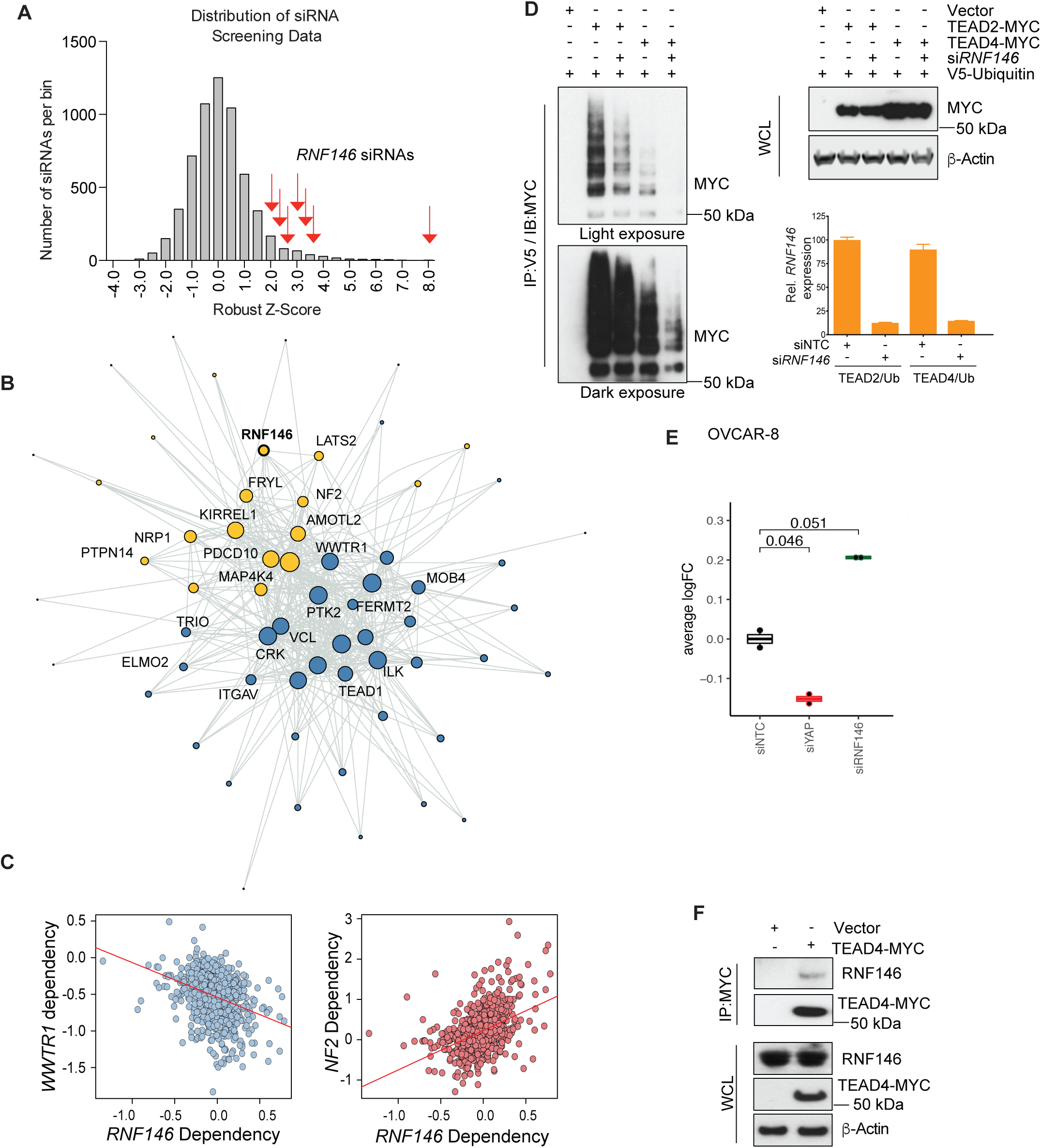
RNF146 is a putative ubiquitin ligase for TEAD. **(A)** RNF146 was identified from an E3 ligase screen using siRNA libraries targeting 1088 ubiquitin-related genes. All seven siRNAs of RNF146 were identified from the screen. The x-axis represents the calculated robust Z-score and the y-axis represents the number of siRNAs per bin. **(B)** RNF146 is a negative regulator of the Hippo pathway. DepMap dependency network of YAP/TAZ. Nodes are significantly correlated genes, significantly correlated genes are connected by a line. Colors of nodes denote 2 distinct clusters based on a random walk algorithm. **(C)** The RNF146 Chronos score is negatively correlated with the Chronos score of *WWTR1* and positively correlated with the Chronos scores of *NF2*. **(D)** TEAD ubiquitination is reduced upon knockdown of *RNF146*. After 24 hours of siRNF146 treatment, TEAD2-Myc or TEAD4-Myc and V5-ubiquitin were transfected into HEK293 cells for an additional 48 hours, and a ubiquitination assay was performed as described in Figure 1. TEAD expression and ubiquitinated TEAD were detected with anti-Myc antibody. RNF146 knockdown was confirmed via Taqman RT-qPCR (n=3). **(E)** RNF146 negatively regulates the Hippo signature. OVCAR-8 cells were transfected with siNTC, si*RNF146*, or si*YAP1* as indicated for 48 hours. RNA-seq was performed, and the Hippo signature score was calculated. **(F)** TEAD interacts with RNF146. HEK293 cells were transfected with TEAD4-Myc for 48 hours. TEAD4 pull-down was then performed with an antibody that recognizes the Myc-tag. TEAD4 expression and RNF146 interaction were detected using Myc antibody and RNF146 antibody, respectively.

RNF146 had been previously reported to enhance YAP activity through the degradation of AMOTs (Wang et al., 2015). To understand this discrepancy with our finding, we leveraged whole-genome dependency screen results from DepMap to broadly assess the loss-of-function effect of RNF146 across cancer cell lines (Meyers et al., 2017). We systematically mapped the functional interactome of the Hippo pathway by assessing the correlation between the loss-of-function effect of individual genes (measured by their Chronos scores) with that of the known Hippo pathway components across 739 cancer cell lines. As expected, we captured strong positive correlations between *WWTR1/TAZ* versus known negative regulators of the Hippo pathway, including *NF2*, *LATS2*, *AMOTL2*, and *MAP4K*. Intriguingly, we found that the *RNF146* Chronos score is anti-correlated with the Chronos scores of TEAD1 and WWTR1/TAZ similar to those known negative regulators (Figure 2B and 2C). Taken together, this provided functional evidence for E3 ligase RNF146 as a putative negative regulator of the Hippo pathway transcription effectors in YAP/TAZ-dependent cancer cell lines.

To further assess the function of RNF146, we depleted RNF146 by either siRNA-mediated knockdown (Figure 2D) or CRISPR-Cas9 knockout (Figure S2C). We observed a significant reduction of TEAD2 and TEAD4 polyubiquitination (Figure 2D and S2C). Additionally, we performed RNA-seq and examined the effects of *RNF146* knockdown on the transcriptional output by YAP/TAZ-TEAD using a published Hippo signature (Pham et al., 2021). We found that si*RNF146* led to the up-regulation of Hippo signature genes, while siY*AP1* led to the down-regulation of the Hippo signature (Figure 2E and S2D, E, and F). This is consistent with the role of *YAP1* as a transcriptional coactivator for TEAD while RNF146 negatively regulates TEADs. We further examined whether RNF146 interacts with TEAD by performing immunoprecipitation using a Myc-tagged TEAD4 construct, and confirmed the interaction between TEAD4 with endogenously expressed RNF146 (Figure 2F).

To assess the effects of RNF146 depletion on endogenously expressed TEADs, we used siRNA and sgRNA to reduce RNF146 level, and performed a cycloheximide chase for 15 hours to monitor TEAD1 and TEAD4 stability in four different cancer cell lines. RNF146 depletion stabilized both TEAD1 and TEAD4 in all four cell lines (Figure S2G-I), although the effect on TEAD1 was less pronounced. Such differential effects on TEAD1 and TEAD4 are consistent with their differential levels of ubiquitination as shown above (Figure 1A). In addition, we performed a cycloheximide chase time course to compare the stability of TEAD1 and TEAD4 subject to siRNF-146 and proteasome inhibitor treatment in OVCAR-8 cells (Figure S2G). Indeed, TEAD1 appears more stable than TEAD4, and RNF146 depletion or proteasome inhibition stabilized both TEAD paralogs. These results indicate that while RNF146-mediated proteasome degradation is a common mechanism that can reduce the stability of different TEADs, some TEAD paralogs such as TEAD1 are more labile. These results are consistent with the notion that the TEAD family of transcription factors is generally quite stable in cells.

Lastly, we performed a proximity ligation assay (PLA) to confirm RNF146 and full-length nuclear TEAD4 are present in a complex using antibodies specific to each protein (Figure S3A-D). We observed nuclear puncta in PA-TU-8902 cells, suggesting the interaction between full-length TEAD4 and RNF146. Taken together, our studies demonstrate that RNF146 interacts with TEAD4, induces TEAD4 polyubiquitination, and negatively regulates TEAD-mediated gene transcription.

#### The ubiquitination of TEAD is dependent on PARylation

*RNF146* has been reported to be a poly (ADP-ribosyl)ation (PARylation)-dependent E3 ubiquitin ligase (Kang et al., 2011; Zhang et al., 2011; Zhou et al., 2011). Therefore, we examined whether TEAD4 ubiquitination was also dependent on PARylation. IP-mass spectrometry studies with endogenous TEAD4 identified two members of poly (ADP-ribose) polymerases, including PARP1 and PARP9 (Figure S3E and Supplemental Table 3). As PARP9 is a mono-ADP-ribosylation enzyme and only PARylates ubiquitin, we focused on PARP1 (Calses et al., 2023; Yang et al., 2017). First, we confirmed the interaction between TEAD4 and PARP1 by IP and Western blot (Figure 3A). Next, we transfected TEAD4 expression constructs into HEK293 cells and blotted immunoprecipitated TEAD4 with an anti-PAR antibody to confirm its PARylation (Figure 3B). Furthermore, we demonstrated that TEAD4 ubiquitination partially depends on PARP1, as *PARP1* depletion using sgRNA reduced TEAD4 ubiquitination levels (Figure 3C and S3F). While these results indicate PARP1 is likely responsible for TEAD4 PARylation and subsequent ubiquitination, the incomplete loss of TEAD4 ubiquitination suggests potential roles of other PARPs and PARylation-independent mechanisms of ubiquitination in the process.

**Figure 3.**
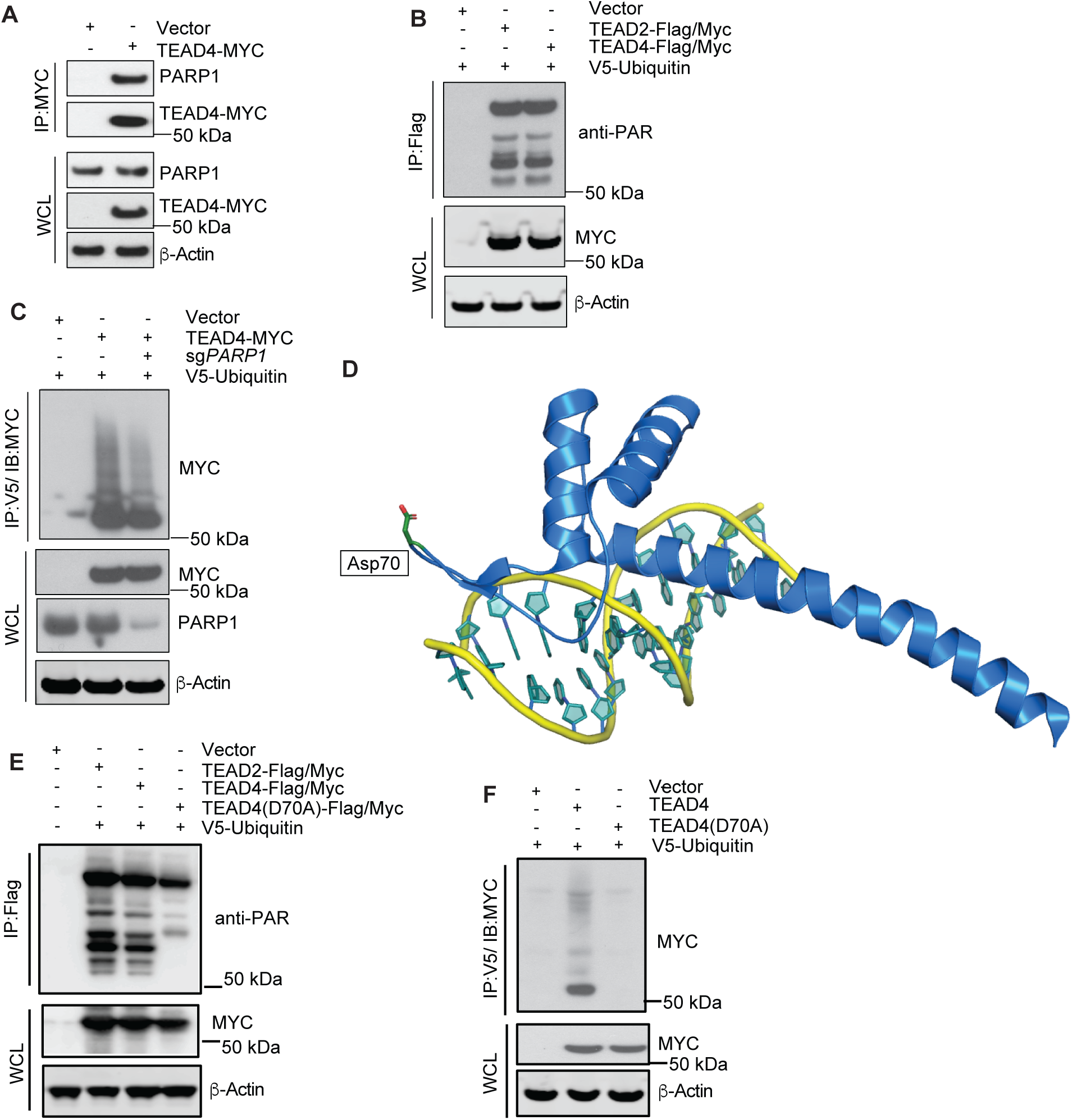
The ubiquitination of TEADs is dependent on their PARylation state. **(A)** TEAD4 interacts with PARP1. HEK293 cells were transfected with TEAD4-Myc plasmid for 48 hours, followed by immunoprecipitation with anti-Myc antibody. PARP1 expression was detected with an anti-PARP1 antibody (n=3). **(B)** TEAD2 and TEAD4 are PARylated. HEK293 cells were transfected with duo-tagged TEAD2-Myc-Flag or TEAD4-Myc-Flag and V5-ubiquitin plasmids for 48 hours, followed by immunoprecipitation with anti-Flag antibody. TEAD expression and PARylation were detected with anti-Myc and anti-PAR antibodies, respectively (n=3). **(C)** TEAD4 ubiquitination is significantly reduced upon knockdown of *PARP1*. HEK293 cells were transfected with specific guide RNAs to knock down *PARP1*. The control and *PARP1* knockdown cell lines were transfected with TEAD4-Myc and V5-ubiquitin plasmids for 48 hours, followed by immunoprecipitation with anti-V5 antibody. TEAD4 expression and ubiquitinated TEAD4 were detected with anti-Myc antibody (n=3). **(D)** PARylation of TEAD is located in the DNA binding domain (Asp70 site on TEAD4). Structure of TEAD4 DNA binding domain in complex with DNA (Shi et al., 2017) with labeled sites for PARylation (green). **(E)** TEAD4(D70A) mutation affects PARylation. HEK293 cells were transfected with duo-tagged TEAD2-Myc-Flag, TEAD4-Myc-FLAG or TEAD4(D70A)-Myc-FLAG and V5-ubiquitin plasmids for 48 hours, followed by immunoprecipitation with anti-Flag antibody. TEAD expression and PARylation were detected with anti-Myc and anti-PAR antibodies, respectively (n=2). **(F)** The ubiquitination of TEAD is significantly reduced in the absence of PARylation. HEK293 cells were transfected with TEAD4-Myc or TEAD4(D70A)-Myc and V5-ubiquitin plasmids for 48 hours, followed by immunoprecipitation with an anti-V5 antibody. TEAD4 expression and ubiquitinated TEAD4 were detected with anti-Myc antibody (n=3).

PARylation has been reported to play diverse roles in the DNA damage response (DDR), DNA repair, transcription, replication, chromatin remodeling, metabolism, and cell death (Kamaletdinova et al., 2019). To pinpoint the PARylation site on TEADs and further dissect the relationship between TEAD PARylation and ubiquitination, we leveraged previous proteome-wide mass spectrometric studies for PARylation (Jungmichel et al., 2013; Zhang et al., 2013) and identified PARylation of a conserved aspartic acid (Asp70) site on TEAD 1-4 in the TEAD DNA binding domain (Shi et al., 2017). Notably, this site is distal from the DNA binding surface and lies in the linker region between two α-helices that interact with the DNA (Figure 3D). Intriguingly, mutating Asp70 to an alanine (Ala) significantly decreased TEAD4 PARylation (Figure 3E) and ubiquitination (Figure 3F), and extended TEAD4 half-life (Figure S3G). These data suggest that PARylation by PARP-family proteins marks TEAD4 for ubiquitination by PARylation-dependent E3 ubiquitin ligase RNF146.

### RNF146 is a negative regulator of the Hippo pathway in *Drosophila melanogaster*

The Hippo pathway is conserved in *Drosophila melanogaster* and mammals (Halder and Johnson, 2011; Pan, 2010; Sudol and Harvey, 2010; Zhao et al., 2010). Given the sequence similarity of *Drosophila* Scalloped (Sd) and mammalian TEAD1-4, including the conserved Asp70 residue (Figure 4A), we examined the genetic interaction between *RNF146* and the Hippo pathway in *Drosophila* using the tissue overgrowth phenotype of Hpo loss-of-function conditions (Udan et al., 2003).

**Figure 4.**
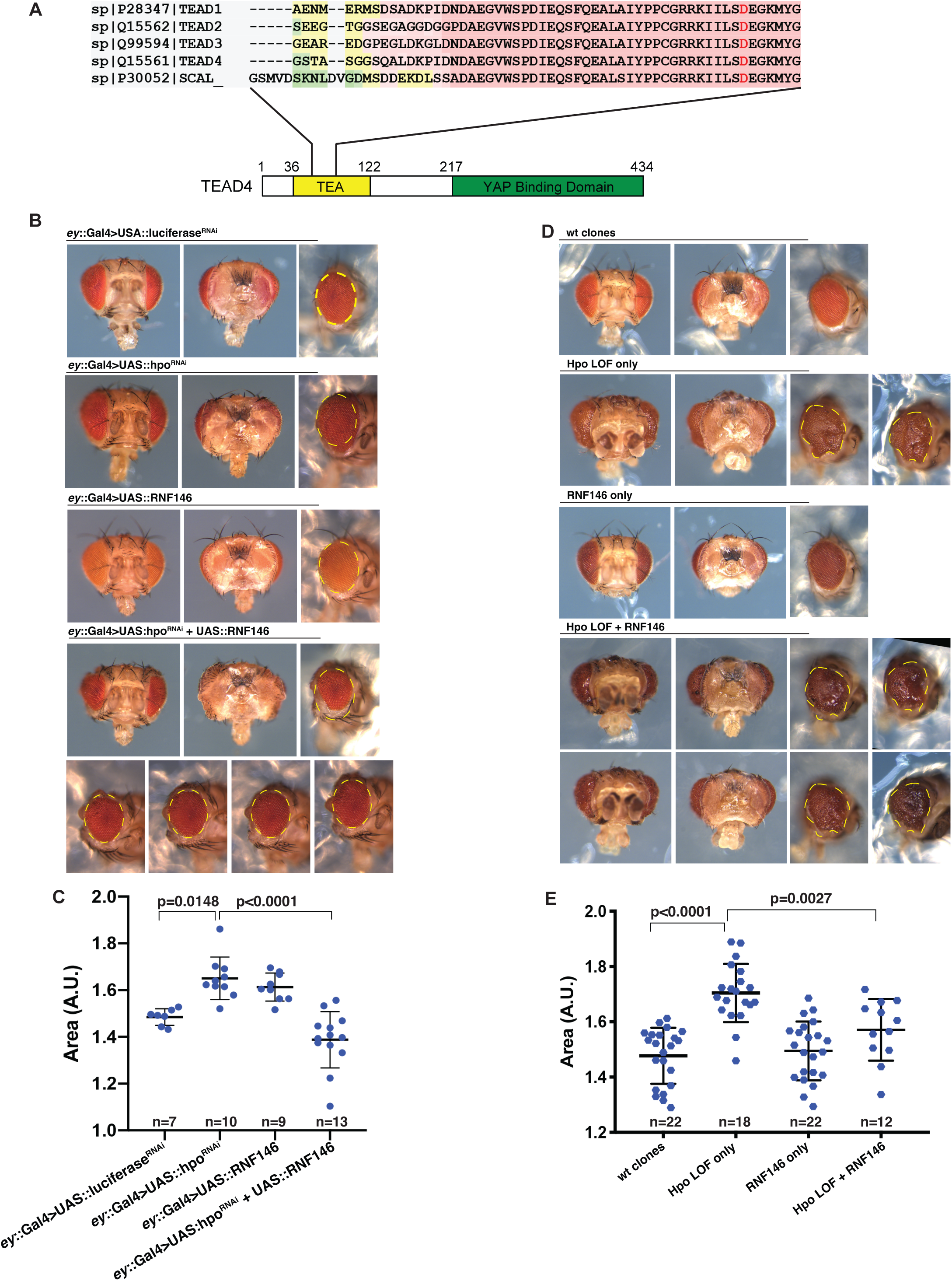
Overexpression of RNF146 E3 ligase antagonizes the overgrowth phenotype caused by the Hpo mutant. **(A)** PARylation site for TEAD is conserved in humans and *Drosophila melanogaster*, as shown in the alignment of TEAD1/2/3/4 and Sd protein sequences. **(B, C)** Overexpression of RNF146 can rescue the overgrowth phenotype caused by Hpo RNAi. B, the first group shows the flies with a control RNAi (ey:: Gal4>UAS:: luciferase RNAi) (n=7). The second group shows knockdown of Hpo (w1118; ey::Gal4/+: UAS::hpoRNAi/+) resulted in a significantly larger eye (n=10). The third group shows flies over expressing RNF146 (ey::Gal4>UAS::RNF146) (n=9). The fourth group, overexpression of RNF146 in a hyperactive Yki background (Hpo RNAi) rescues the overgrowth phenotype resulting in a significantly smaller eye (n=13). C is the quantification of B. Means +/- S.D. of eye area quantified per genotype are shown in arbitrary units (A.U.). One-way ANOVA with Sidak’s multiple comparisons. Numbers (n) of eye area quantified and P-values are indicated. **(D, E)** Overexpression of RNF146 can rescue the overgrowth phenotype caused by a Hpo loss of function mutation. D, Wild-type (wt) control clones were generated using the eyFLP system (genotype: w1118; neoFRT42D, +/neoFRT42D, GMRHid; ey::Gal4, UAS::FLP/+) (Newsome et al., 2000) (n=11, both eyes). *hpo^KC202^* mutant clones (genotype: *w^1118^*; FRT42D, hpoKC202/FRT42D, GMRHid; ey::Gal4,UAS::FLP/+) result in a large, folded eye with cuticle overgrowth around the eye. This increase in eye area was significant compared to the eyes of WT clones. (p<0.0001; n=9, both eyes). Clones overexpressing RNF146 showed no phenotype compared to control (genotype: *w^1118^*; FRT42D, + /FRT42D, GMRHid; *ey*Gal4, UAS::FLP/UAS::RNF146^OE^; n=11, both eyes). Combining *hpo^KC202^* and RNF146 overexpression (genotype: *w^1118^*; FRT42D, hpoKC202/FRT42D, GMRHid; ey::Gal4,UAS::FLP/ UAS::RNF146^OE^) resulted in a significantly smaller eye as compared to the *hpo^KC202^* mutant only. (p=0.0027; n=6, both eyes). E is the quantification of the eye area of all eyes in D. Means +/- S.D. of eye area quantified per genotype are shown in arbitrary units (A.U.). One-way ANOVA with Sidak’s multiple comparisons. Numbers (n) of eye areas quantified and P-values are indicated.

First, we induced Hpo loss-of-function in the developing eye by expressing a dsRNA against Hpo using ey::Gal4. This resulted in significantly larger eyes with additional folding of the cuticle behind the eyes compared to luciferase RNAi as a control. We confirmed that Sd is required for eye overgrowth in Hpo loss-of-function conditions, as Hpo knockdown did not increase eye size in Sd RNAi-expressing animals (Figure S4A, B). These phenotypes are consistent with inactivation of the Hippo pathway kinase cascade resulting in nuclear localization of Yki, which then binds Sd to activate cell proliferation. In such a Hpo loss-of-function background, we found that overexpression of RNF146 rescued the overgrowth phenotype and significantly reduced eye size, supporting its role as a negative regulator of the transcriptional program downstream of Sd (Figure 4B, C). Conversely, RNF146 knockdown increased eye size further in animals expressing Hpo RNAi, indicating that RNF146 can negatively regulate the activity of Sd/Yki and limit their full activation in Hpo loss of function conditions (Figure S4C, D). Next, we assessed the genetic interaction between RNF146 and Hpo using an eyFLP system (Newsome et al., 2000) to generate *hpo^KC202^* mutant clones and overexpress RNF146 in a genetic background in which non-clonal cells (twinspots) are eliminated by the presence of GMR::Hid (Newsome et al., 2000). Using this system, we again observed that overexpression of RNF-146 suppresses the overgrowth phenotype of Hpo mutant clones compared to wild-type clones (Figure 4D, E). Of note, overexpression of RNF146 alone resulted in a modest increase in eye size compared to luciferase RNAi control (Figure 4C). This is in contrast to the anti-growth effect of RNF-146 in the Hpo loss-of-function background and indicates RNF146 may regulate other genes/pathways controlling eye sizes besides its role as a negative regulator of Sd/yki activity. Taken together, the genetic interaction between RNF146 and Hpo in *Drosophila* eyes demonstrates that RNF146 functions as a negative regulator of the over-growth phenotype associated with dysregulation of the Hippo pathway, consistent with our results in human cell line models described above.

#### TEAD1/2/3/4 down-regulation suppresses cell proliferation and Hippo pathway signaling

Deregulation of Hippo signaling pathway has been implicated in various human cancers. To understand the effects of TEAD loss, we performed *TEAD1-4* knockdown to determine TEAD dependency across a spectrum of cancer cell lines. We confirmed *TEAD1-4* knockdown via Taqman RT-qPCR and Western blotting three days after transfection (Figure S5A and S5B) and assessed cell proliferation using the Cell Titer Glo assay. *TEAD1-4* knockdown downregulated the Hippo pathway signature gene *CTGF* (Figure S5A) suggesting the pathway was altered. *TEAD1-4* knockdown significantly reduced cell proliferation in all cell lines tested except two negative control Hippo-independent cell lines SK-N-FI and HCSC-1, which lack *YAP/WWRT1* expression (Figure 5A). We further confirmed TEAD dependency *in vivo* using xenograft studies in mice. Mice (n=10) injected with MDA-MB-231 cells showed a significant reduction in tumor volume upon induced *TEAD1-4* knockdown in an intervention setting (Figure 5B and S5C-D). Together, these results indicate the depletion of TEADs could serve as a therapeutic strategy to inhibit growth of Hippo-pathway altered cell lines. While we identified RNF146 as an E3 ligase for TEAD transcription factors, TEAD transcription factors are generally very stable (Pham et al., 2021). We reasoned that their slow turn-over rate, the intermediate step of being PARylated, as well as different ubiquitination levels of all four TEAD paralogs (Figure 1A and S2G), may limit the ability of RNF146 and potentially other endogenous E3-ligase to reduce TEAD protein levels in cells. To overcome the limitations of the endogenous mechanisms of TEAD degradation, we went on to develop heterobifunctional CIDEs to pharmacologically induce TEAD degradation.

**Figure 5.**
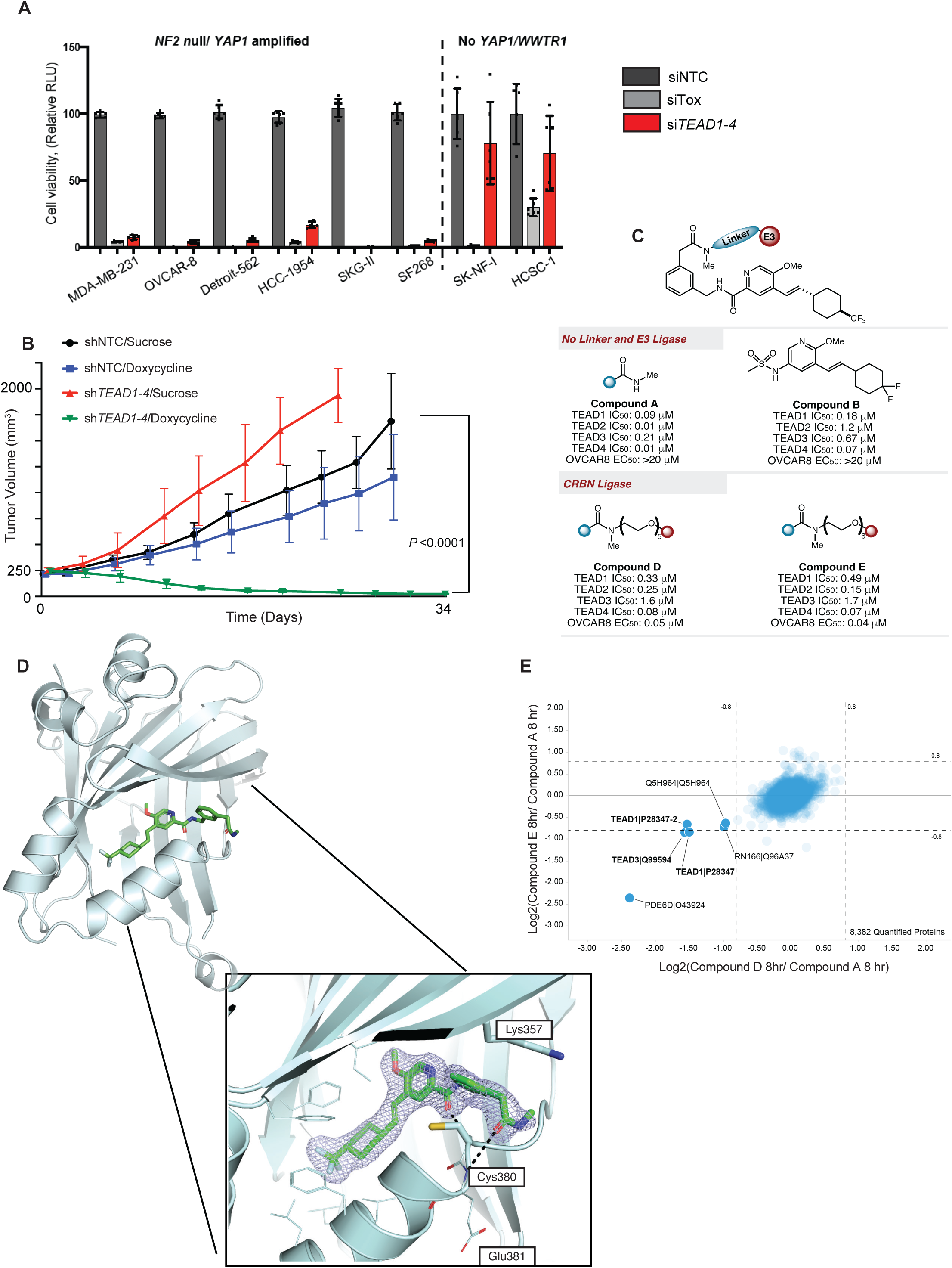
*TEAD* knockdown induces robust Hippo pathway suppression. **(A)** Cell viability was assessed by Cell Titer-Glo upon siRNA treatment of *TEAD1-4* in Hippo/TEAD sensitive (*NF2* null/*YAP1* amplified) and insensitive cell lines (no YAP1/WWTR1). Cells were treated with siNTC (dark gray bars), siTox (light gray bars) and siTEAD1-4 (red bars) for 72 hours before the cell viability assay. **(B)** *In vivo* xenograft growth (measured as tumor volume over time) of MDA-MB-231 cells expressing TEAD1-4 shRNA in the presence of sucrose (red) or doxycycline (green), or expressing shNon-Targeting Control (NTC) in the presence of sucrose (black) or doxycycline (blue). **(C)** Chemical structures, biochemical and cellular activities of all CIDEs screened. **(D)** Crystal structure of Compound A bound to TEAD. **(E)** Protein abundance measurements were made using tandem mass tag quantitative mass spectrometry and plotted as a log fold change. The scatterplot depicts the change in relative protein abundance in MDA-MB-231 cells treated with Compound D (x-axis) and Compound E (y-axis) relative to Compound A. Cells were treated with 0.5 µM of the compounds for 8 hours before harvesting.

#### Development and Screening of pan-TEAD lipid pocket degraders

We designed and synthesized a series of heterobifunctional TEAD CIDEs based on reversible TEAD lipid pocket binding ligands A and B. Compound A is a close-in analog to Compound B, as described previously (Holden et al., 2020). Compound B binds the same site as Compound A in a similar manner, and the only modifications to Compound B were inclusion of a handle for attaching an E3 Ligase ligand. These compounds have been previously disclosed in patent application WO2020051099. To facilitate the structure-based design of potent TEAD degraders, we solved the co-crystal structure of Compound A bound to TEAD2 (Figure 5C, D). As expected, based on previous reports and our biochemical data, this compound strongly binds all the TEAD isoforms via the lipid pocket in a TEAD lipid HTRF assay. Compound A makes hydrogen bonds to the backbone amides of Cys380 and Glu381. Still, binding appears to be driven primarily by van der Waals interactions deep in the pocket and with the benzyl group, sandwiched between Cys380 (which would otherwise be lipid-modified) and Lys357 (Figure 5D). This moiety provides an optimal vector for the terminal amide to exit the pocket, allowing for the conjugation of an E3 ligase ligand via a linker in our TEAD-CIDEs (Figure 5D and S5E). A set of TEAD CIDEs containing ligands that bind the ubiquitin ligases XIAP, CRL2-VHL, and CRL4-CRBN with various linker lengths was generated and evaluated. Reversible TEAD lipid pocket ligands, Compound A and Compound B were used as non-degrader control compounds (Figure 5C). Gratifyingly, TEAD-CIDEs based on Compound A maintained good pan-TEAD biochemical potency when linked to all three evaluated ligase ligands. TEAD/CRBN CIDEs also led to significant activity in our cellular OVCAR-8 antiproliferation assay (Figure 5C). Compound D and Compound E were selected for further evaluation based on these data. Attempts to obtain co-crystal structures of TEAD, CRBN, and either of the compounds were unsuccessful, likely owing to the highly flexible nature of the PEG linkers in our compounds.

We next assessed Compound D and Compound E for TEAD specificity and selectivity using multiplexed mass spectrometry-based global proteomics in the MDA-MB-231 cell line. We identified significant downregulation of shared and unique TEAD peptides (TEAD1, 3) when cells were treated with Compound D and Compound E for both 4 hours and 8 hours compared to Compound A (Figure 5E and S5F). Other than TEAD peptides, we noted that PDE6δ (Retinal rod rhodopsin-sensitive cGMP 3’,5’-cyclic phosphodiesterase subunit delta) and RNF166 (Ring finger protein 166) were the only other significantly reduced proteins (Figure 5E and S5F). RNF166 has been previously identified as a target of other CRBN-based ligands (lenalidomide) and degraders (You et al., 2020), indicating it could be an off-target for the CRBN binding portion of Compound D and Compound E. As expected, the parent ligand Compound A did not induce any significant changes in protein abundance, indicating TEAD degradation induced by CIDEs depends on E3 ligase recruitment.

#### CIDE-induced TEAD degradation is dependent on CRBN modulation and the proteasome

We next evaluated and confirmed that Compound D and Compound E induced TEAD degradation in TEAD-dependent cell lines in a concentration- and time-dependent manner by Western blot (Figure 6A, 6B, S6A). No TEAD degradation was observed upon treatment with a TEAD lipid pocket binder (compound A or compound B) (Figure 6A and S6B).

**Figure 6.**
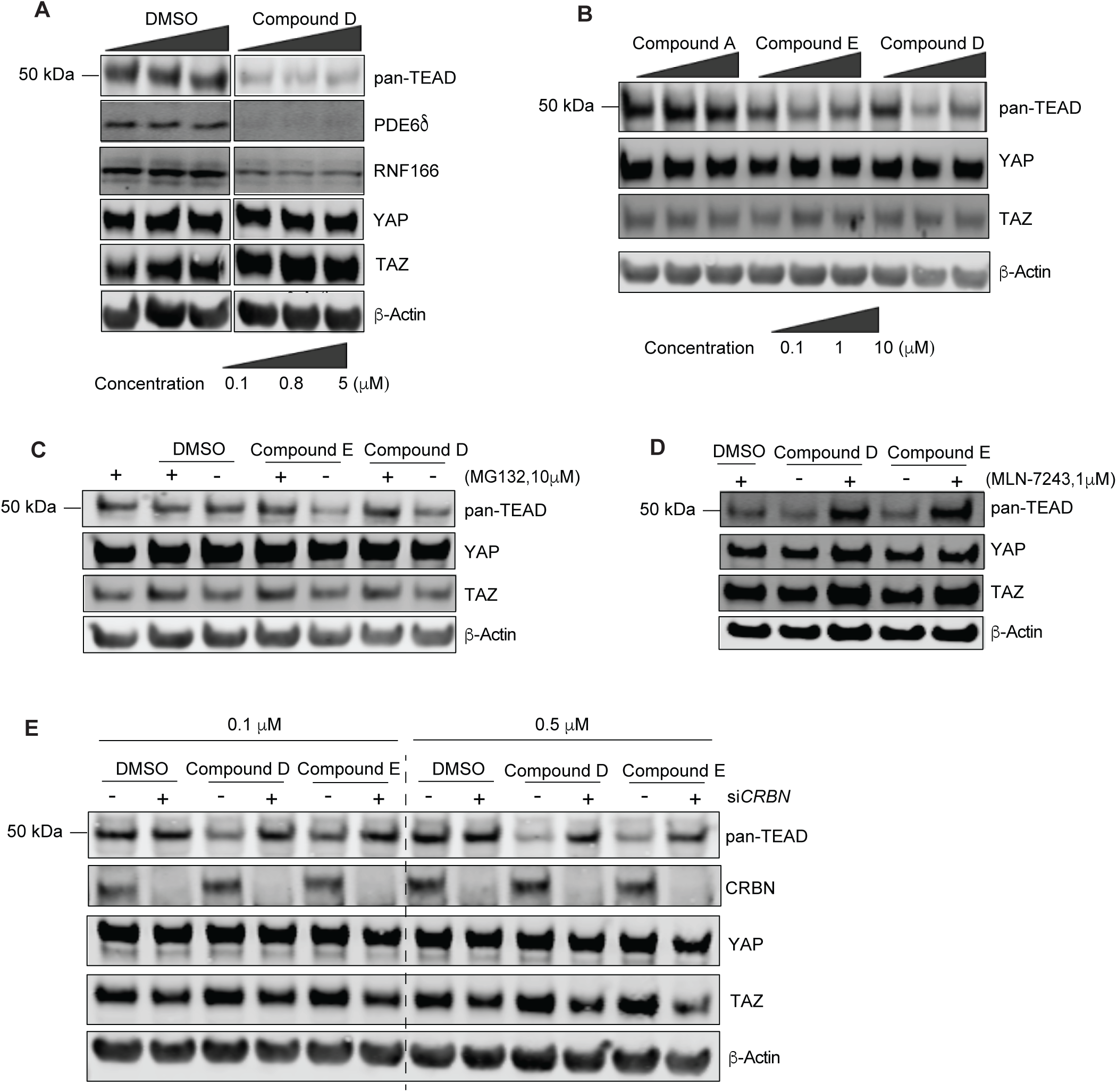
CIDE-mediated TEAD degradation is dependent on CRBN and the ubiquitin-proteasome pathway. **(A)** Immunoblot for pan-TEAD, PDE68, RNF166, YAP, TAZ, and β-Actin in MDA-MB-231 cells after 24 hours of Compound D treatment at the indicated concentrations (0.1 µM, 0.8 µM, 5 µM). **(B)** Immunoblot for pan-TEAD, YAP, TAZ, and β-Actin in OVCAR-8 cells after 16 hours of treatment with Compound A/Compound E/Compound D at the indicated concentrations of 0.1 μM, 1 µM and 10 µM. **(C)** Immunoblot for pan-TEAD, YAP, TAZ, and β-Actin in OVCAR-8 cells upon co-treatment with DMSO/Compound E /Compound D (1 µM) for 5 hours and MG132 (10 µM) for 6 hours. **(D)** Immunoblot for pan-TEAD, YAP, TAZ, and β-Actin in MDA-MB-231 cells upon co-treatment with DMSO/Compound D/Compound E (1 µM) for 5 hours and MLN-7243 (1 µM) for 6 hours. **(E)** Immunoblot for pan-TEAD, CRBN, YAP, TAZ, and β-Actin in MDA-MB-231 cells upon treatment with DMSO/Compound E/Compound D (0.1 µM and 0.5 µM) for 16 hours. siRNA was transfected 3 days prior to compound treatment as indicated.

This confirms the mass-spec quantifications and indicates that TEAD and CRBN engagement are required for CIDE-mediated TEAD degradation. In addition, we also assessed PDE6δ and RNF166 abundance and confirmed their degradation by Compound D (Figure 6A). Based on this result, we docked Compound A into the crystal structure of the PDE6δ ligand binding pocket (PDB ID: 5ML2) and found that it could occupy that pocket without significant clashes (Figure S6C) (Tian et al., 2010). However, from a genome-wide CRISPR screen (Meyers et al., 2017), PDE6δ dependency was not correlated with other Hippo pathway members (Figure S6D, E). It was not essential in the TEAD-dependent cell lines tested here, suggesting that PDE6δ loss would not lead to anti-growth effects in the TEAD-dependent cell lines used in our study.

Next, we treated cells with proteasome and ubiquitin ligase E1 inhibitors. Co-treatment of cells with Compound D/Compound E with MG132 (a proteasome inhibitor) or MLN7243 (an E1 inhibitor) prevented TEAD degradation (Fig 6C, 6D), indicating that the proteasome pathway is required for effective CIDE-mediated degradation. To confirm that Compound D and Compound E-induced TEAD degradation is CRBN-dependent, we performed siRNA knockdown of *CRBN* in MDA-MB-231 and observed rescue of TEAD degradation induced by Compound D (Figure 6E). Overall, the data shows that TEAD-CIDEs-mediated TEAD degradation depends on the ubiquitin/proteasome pathway and the CRL4-CRBN ubiquitin ligase.

#### TEAD-CIDEs inhibit proliferation and downstream signaling in Hippo pathway-dependent cell lines

To assess the anti-proliferative effect of TEAD-CIDEs, we tested Compound D and Compound E in OVCAR-8 and SK-N-FI (*YAP/WWTR1* deficient, Hippo pathway independent) cells. Both TEAD-CIDEs decreased cell proliferation in the OVCAR-8 cell line and did not have anti-proliferative effects in SK-N-FI (Figure 7A, 7B). No anti-proliferative effect was observed with Compound A or B, suggesting that this effect was degradation-dependent (Figure 7A, 7B). Similar effects were observed in a colony formation assay (Figure 7C, 7D). The anti-proliferative effect of Compound D was further validated across a spectrum of Hippo pathway-dependent cell lines (Figure S7A, S7B). Given the enhanced anti-proliferative effects of Compound D compared to Compound A, we next evaluated its effects on Hippo pathway output by RNA-seq. Compound D treatment resulted in robust concentration-dependent downregulation of Hippo pathway target genes compared to Compound A (Figure 7E and 7F). Taken together, our data serves as a proof-of-concept that TEAD proteins can be pharmacologically targeted and modulated through the ubiquitin-proteasomal system.

**Figure 7.**
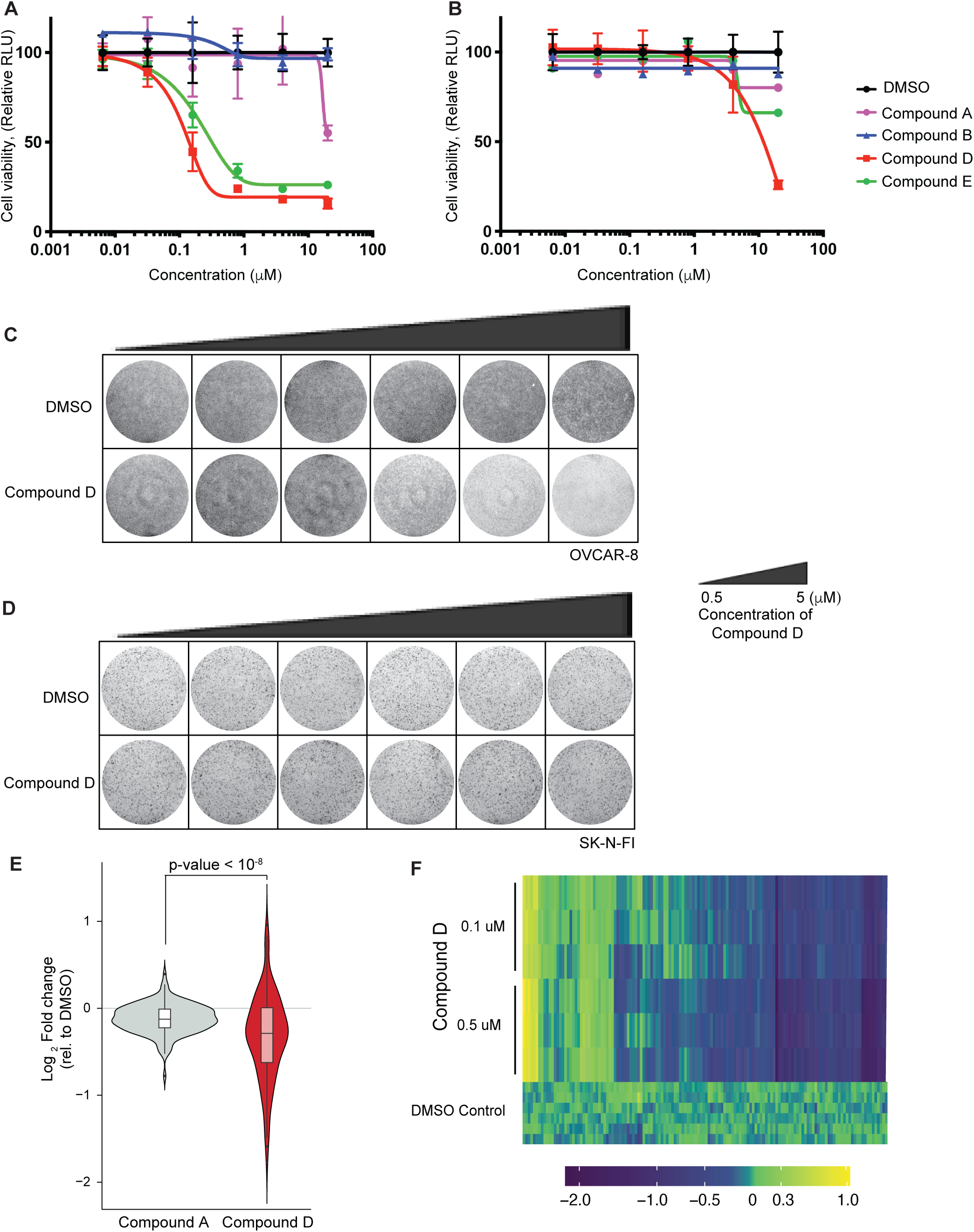
TEAD degradation inhibits cell proliferation and downstream Hippo pathway signaling. **(A, B)** Cell viability was assessed using CellTiter-Glo in (A) OVCAR-8 cells, or in (B) SK-N-FI cells after 6 days of treatment with Compound D (red), Compound E (green), Compound A (purple), Compound B (blue), or DMSO (black). Error bars represent the standard deviation of five technical replicates. **(C, D)** Crystal violet growth assay in OVCAR-8 cells (Hippo dependent) and SK-N-FI cells (Hippo independent) after 6 days treatment with Compound D at indicated concentrations. **(E, F)** RNA-seq of Hippo signature genes after Compound D/Compound A treatment in MDA-MB-231 cells. Hippo signature genes exhibit significant downregulation in a concentration dependent manner upon treatment with Compound D. Cells were treated with 0.1 µM and 0.5 µM of Compound A or Compound D for 24 hours before harvesting.

#### TEAD CIDE specifically reduced chromatin accessibility at TEAD motifs

To assess whether TEAD-CIDE could alter the chromatin landscape in cells, we performed ATAC-seq on OVCAR-8 cells treated with Compound D or DMSO control for 48 hours. Differential peak analysis detected 4394 gained, 9482 lost, and 132803 unaltered regions (absolute logFC >1 and FDR < 0.01, negative binomial test) (Figure 8A). Both the gained and lost regions were predominantly distal to protein coding genes (Figure 8B). We observed significant reductions in chromatin accessibility at the promoter and multiple enhancer regions of well-known TEAD targets such as ANKRD1 and CCN1 (a.k.a. CYR61) (Figure 8C).

**Figure 8.**
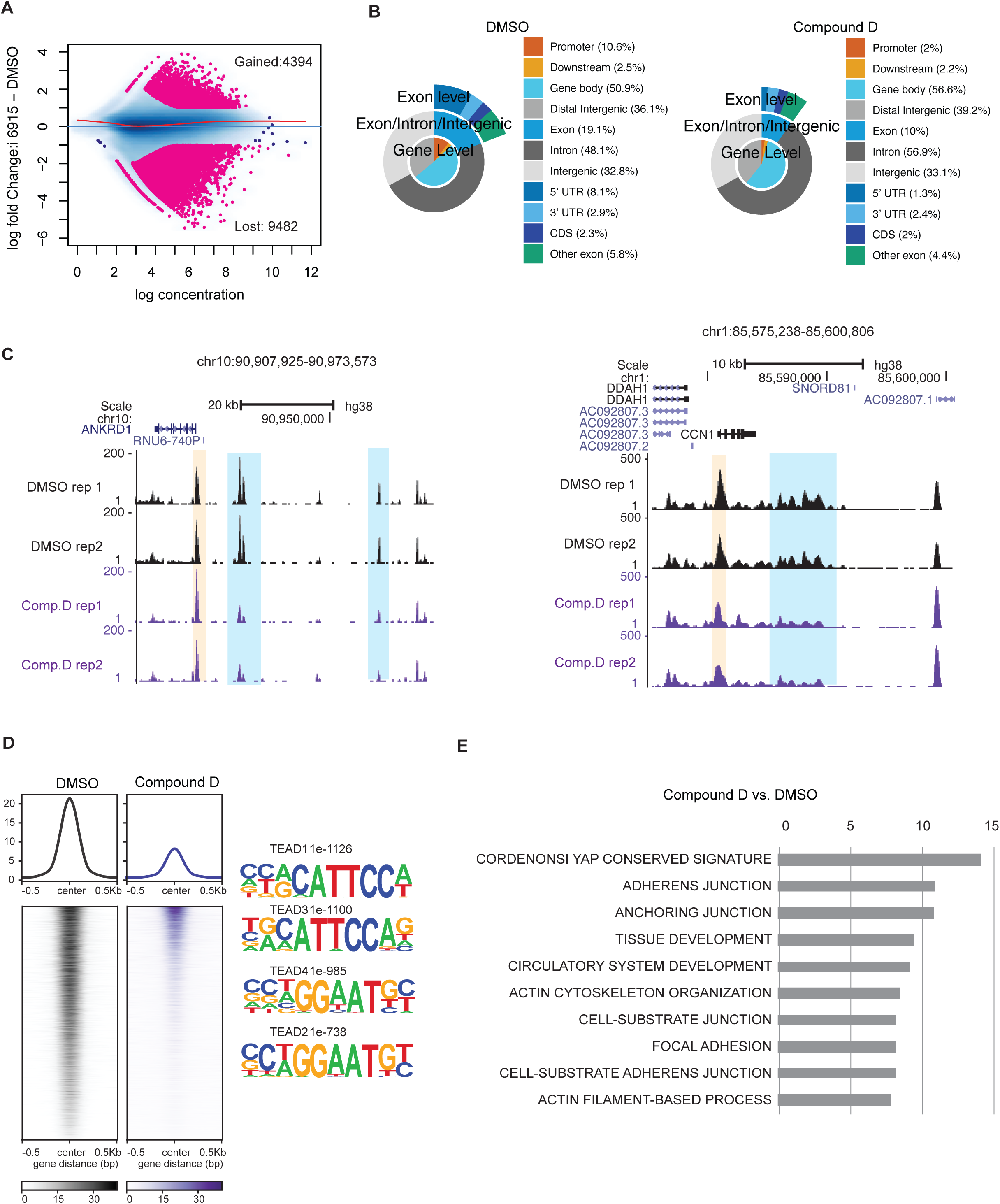
Compound D specifically reduces chromatin accessibility at TEAD motifs. **(A)** Differential analysis of ATAC-seq of OVCAR-8 cells treated with DMSO or Compound D for 48 hours using DiffBind. **(B)** Genomic annotations of the differential ATAC-seq peaks by ChIP-Annotate. **(C)** UCSC browser view of ATAC-seq at known targets of TEAD including ANKRD1 and CCN1/CYR61. **(D)** Lost ATAC-seq peaks were visualized by Deeptools and analyzed for motif enrichment by HOMER. **(E)** Lost peaks were assigned to genes using known distal enhancer-gene target links in Poly-Enrich, and the 10 most significant pathways are shown.

Systematic unbiased motif enrichment analysis revealed that regions associated with loss of chromatin accessibility were significantly enriched for TEAD DNA binding motifs (Figure 8D). Since most downregulated peak regions were distal, we assigned enhancer-gene targets using links derived from aggregated chromatin capture data by Poly-Enrich. The most significantly enriched pathway corresponded to Cordenonsi_YAP_conserved_signature (Figure 8E). This suggests that the observed downregulation of Hippo pathway target genes is likely due to the loss of TEAD-associated chromatin accessibility *in vitro*. Interestingly, we noted that among the ten most downregulated pathways, many were related to cell adhesion (Figure 8E), which is linked to YAP/TAZ biology (Dey et al., 2020). These results confirm that the TEAD CIDEs can modulate known YAP-TAZ regulated cellular processes and exert a TEAD degradation-dependent anti-growth effect on cancer cells. Collectively, our data highlights TEAD degraders as valuable tools to uncover the role of TEAD accessibility in cancers to guide future drug development efforts.

## DISCUSSION

In this study, we show that the activity of TEAD transcription factors can be post-transcriptionally regulated via the ubiquitin/proteasome system. We demonstrate that TEAD proteins can be poly-ubiquitinated (Figure 1) and identify RNF146 as an E3 ligase for TEADs (Figure 2). Such ubiquitination is dependent, at least in part, on TEAD PARylation by PARP family members (Figure 3). Our study also demonstrates the genetic interaction of RNF146 with the Hippo pathway in *Drosophila melanogaster*, establishing RNF146 as a conserved negative regulator of the Hippo pathway across species (Figure 4). In addition, we observe that RNF146 overexpression induces an overgrowth phenotype on its own. This implies a role of RNF146 in developmental homeostasis in general, which will require future investigation.

RNF146 has been previously reported as a PARylation-dependent ubiquitin ligase (Kang et al., 2011; Zhang et al., 2011; Zhou et al., 2011). TEAD transcription factors have been previously reported as targets of PARylation, which in turn impacts their DNA binding activity (Butler and Ordahl, 1999; Jungmichel et al., 2013; Zhang et al., 2013). The conserved Asp70 PARylation-target residue, and other Asp residues in the DNA-binding domain do not reside in the DNA-binding interface of TEAD4, whereas ubiquitinated Lysine residues do. Thus, we speculate that PARylation of TEAD may indirectly impact DNA-binding through ubiquitination rather than directly affecting DNA-binding. This raises the possibility that ubiquitination by RNF146 may directly modulate TEAD activity in addition to marking them for proteasome-mediated degradation. However, the requirement of PARylation as an intermediate post-translational modification between RNF146 and TEAD, the exact interaction interface between RNF146 and TEAD, and the direct versus indirect role of ubiquitination in regulating TEAD transcriptional activity will need to be further elucidated in future studies.

Previous studies demonstrated that RNF146 may positively regulate YAP activity through degradation of AMOTs (Wang et al., 2015). However, our study demonstrates that RNF146 functions as an E3 ligase and negative regulator of TEADs. We speculate that the different modes of RNF146 action could be influenced by the cellular context, its subcellular localization, and the PARylation status of different RNF146 substrates. Furthermore, our analysis of pan-cancer gene dependency revealed an apparent function of RNF146 as a negative YAP/TAZ/TEAD regulator across many cancer cell lines of diverse indications (Figure 2B, and 2C).

TEADs are central components of the transcriptional machinery controlled by the Hippo pathway. Genetic depletion of TEADs induces strong anti-proliferative effects in Hippo-dependent cancer cell lines. While we identified RNF146 as an E3 ligase for TEAD transcription factors, the knockdown of RNF146 had a limited impact on TEAD stability in cancer cells. Thus, RNF146 may not be the only factor that regulates TEAD stability in cancers, and the knockdown effect of RNF146 may be tissue- and context-specific. This prompted us to pharmacologically engage other ubiquitin ligases and the proteasome system to suppress the activity of all TEAD isoforms and their target genes. To this end, we designed, synthesized, and evaluated a series of potent heterobifunctional TEAD-CIDEs based on reversible TEAD lipid pocket binding ligand Compound A. As a proof-of-concept study, we engineered TEAD-CIDEs (Compound D, Compound E) that could reduce TEAD protein levels in a CRBN-dependent manner with concentrations as low as 0.1 µM. CIDE-induced TEAD degradation resulted in a significant concentration- and time-dependent reduction in downstream Hippo target genes in Hippo pathway-dysregulated cells. To our knowledge, this study demonstrated the first class of TEAD degraders that specifically inhibits cell proliferation and suppresses Hippo downstream signaling, validating TEAD degraders as a potential therapeutic modality to target the Hippo pathway.

We identified PDE6δ as an off-target protein for these TEAD-CIDEs. Structural biology studies suggest that while the pockets of TEAD and PDE6δ are quite different, PDE6δ may still be able to accommodate certain classes of TEAD ligands and should be considered as a potential off-target for TEAD lipid pocket-binding molecules (Tian et al., 2010). While PDE6δ loss is unlikely to impact the observed Hippo pathway specificity, next-generation TEAD-CIDEs should be engineered to achieve higher TEAD lipid pocket binding specificity and degradation efficiency. Further development of TEAD-CIDEs is needed to optimize the pharmacological properties to enable *in vivo* testing.

In summary, we have identified an important mechanism of regulating TEAD protein levels by the ubiquitin/proteasome system and demonstrated that this can be pharmacologically enhanced through TEAD-CIDEs. TEAD-CIDEs serve as potent chemical probes for studying the effects of pan-TEAD degradation for potential therapeutic intervention. As many cancer targets lack enzymatic function, CIDEs enable a tractable option to convert inactive ligands to functional degraders. This study provides further proof-of-concept for degraders as a therapeutic strategy for targeting challenging and hard-to-drug pathways in cancer.

## MATERIALS and METHODS

### Cell lines, antibodies, and other reagents

Cell lines used in this study are HEK293, Detroit 562, OVCAR-8, PA-TU-8902, MDA-MB-231, HCC1576, NCI-H226, MS751, NCI-H2373, SNU-423, HCSC-1 and SK-N-FI which were obtained from American Type Culture Collection (ATCC). MCF7 TEAD-Luc was purchased from BPS Bioscience. Cell line authentication was conducted for Short Tandem Repeat (STR) Profiling using the Promega PowerPlex 16 System. This process was performed once when receiving a new cell line and compared to external STR profiles of cell lines (when available) to determine cell line ancestry. Routine authentication was conducted by SNP-based genotyping using Fluidigm multiplexed assays at the Genentech cell line core facility.

Antibodies used in this study are pan-TEAD (13295, CST); YAP(14074, CST); TAZ (70148, CST); MAX (sc-765 and sc- 8011, Santa Cruz); α-Tubulin (3873, CST); β-Actin (3700, CST); cleaved PARP (9541, CST); FRA1(5281, CST); Myc-tag (2278-CST); V5 (680602, Biolegend); RNF146 (HPA027209, Sigma-Aldrich); TEAD4 (Abcam 58310); PARP1 (9532, CST); anti-PAR (AM-80, Sigma-Aldrich); NUP98 (50610, Abcam); DAPI (4083, CST); CRBN (11435-1-AP, Proteintech); Anti-rabbit and anti-mouse HRP linked (7074 and 7076, CST), IRDye anti-rabbit and anti-mouse (68070 and 32211, Li-cor). siRNAs were purchased from Dharmacon, including siNTC (D-001810-10), si*RNF146* (L-007080-00), si*PARP1* (L-006656-03). MG132 solution (M7449) was from Sigma-Aldrich. MLN7243 was purchased from Selleckchem. On target plus Human CRBN siRNAs (51185) and non-targeting siRNA pool were purchased from Horizon Discovery.

### *In vivo* ubiquitination assay

Myc-Flag-human TEADs (OriGene) and V5-ubiquitin (generated in-house) plasmids were transfected into HEK293T cells at a ratio of 3:1 with Lipofectamine 3000 (Thermo Fisher Scientific). After 48 hours, cells were lysed in NP-40 buffer (1% NP-40, 120 mM NaCl, 50 mM Tris, pH 7.4, 1 mM EDTA, pH 7.4, 20mM cysteine protease inhibitor N-ethylmaleimide (NEM), protease and phosphatase inhibitors (Roche). Proteins were denatured with 1% SDS at 94°C for 10 mins and then diluted to 0.05% SDS before anti-V5 immunoprecipitation. Immunoprecipitation was at 4°C on a turning rotor overnight. After washing with NP-40 buffer and PBS, ubiquitination proteins were then detected by immunoblotting.

#### Immunoprecipitation and immunoblotting

Cells were lysed in RIPA lysis buffer (89900, Thermo Fisher Scientific) containing protease inhibitor (Roche), and phosphatase inhibitor (Roche). Lysates were prepared by taking supernatants from centrifugation at 12,000 g for 15 min at 4°C. Equivalent amounts of proteins were loaded and separated by SDS-PAGE and then transferred to membranes.

For endogenous co-immunoprecipitation experiments, 1×10^7^ cells were lysed using RIPA buffer and immunoprecipitation with indicated antibody overnight at 4°C. After washing with RIPA buffer, co-immunoprecipitated endogenous proteins were detected by immunoblotting.

#### Subcellular Fractionation

Cells grown on 10 cm dishes (80% confluence) were harvested after washing with cold PBS. Cell pellets were then incubated with 200 µl of Buffer A [10 mM HEPES pH 7.9, 10 mM KCl, 1 mM EDTA, 0.1 mM EGTA, 0.2% NP-40, 10% glycerol, complete-mini (EDTA free) protease inhibitor (Roche), PHOStop phosphatase inhibitor (Roche) at 4°C for 20 min. Cytoplasmic fractions were collected by taking the supernatant after centrifugation for 5 minutes at 2500 × g. To prepare the nuclear fraction, the pellets were then incubated with 50 µl of RIPA buffer (89900, Thermo Fisher Scientific) containing protease and phosphatase (Roche) for 30 min at 4°C. After centrifugation for 12 mins at 16,000 × g, nuclear fractions were collected by retaining the supernatant.

#### siRNA Screening

siRNA screening was performed in MCF7 TEAD reporter cells using two different siRNA libraries (Dharmacon On-Target Plus and Ambion Silencer Select) targeting ubiquitin-related genes. The Dharmacon library comprised 4 siRNAs per gene, whereas the Ambion had 3 siRNA per gene. The screening was conducted in biological replications. Briefly, 2 pmol of siRNA (20 nM final concentration) was spotted to 384 well plates, and 0.08 μL Lipofectamine RNAiMax was added in 20 μL of serum-free EMEM media. This complex was incubated at ambient temperature for 30 minutes before adding 2,000 MCF7 TEAD reporter cells in 20 μL of 20% serum EMEM media. Cells were incubated for 72 hours before assaying by OneGlo luciferase assay reagent (Promega). All plates included 8 wells of negative control siRNA (Ambion Silencer Select Negative Control #2), 8 wells of cell death control (Qiagen AllStars Positiv Control siRNA), and 8 wells of YAP targeting siRNA (Ambion Silencer Select, cat# s20366). Positive controls were used to gauge transfection efficiency and yielded assay z’ factors of 0.5 and 0.3 respectively. After normalization of sample wells to negative control siRNA on each plate, robust z-scores (Chung et al., 2008) were calculated. Replicate values were averaged, and gene-level p-values were computed using RSA (König et al., 2007). Parallel screens were also used to assess effects on viability. All data can be found in Supplemental Table 2. All seven siRNAs tested against RNF146 yielded a significant increase in reporter activity (Fig 2A).

#### siRNA transfection

Cells were seeded at 15000/well on a 6-well plate for 24 hours. The following day, cells were treated with the siRNAs (final concentration 20nM) using Lipofectamine RNAi Max in serum-free RPMI media for 72 hours before collection. siRNAs were purchased from Dharmacon, including siNTC (D-001810-10), si*TEAD1* (J-012603-05), si*TEAD2* (J-012611- 09), si*TEAD3* (L-012604-00), siT*EAD4* (J-019570-09), si*RNF146* (L-007080-00), si*PARP1* (L-006656-03), si*CRBN* (51185). Knockdown was confirmed by Western blot with the following antibodies from Cell Signaling Technology: pan-TEAD (13295, CST); YAP (14074, CST); TAZ (70148, CST); β-Actin (3700, CST).

#### RNA extraction, cDNA synthesis and quantitative RT-PCR

Tumors and cell lines were dissociated and lysed for RNA isolation using an RNAeasy kit (Qiagen). cDNA was prepared by reverse transcription using the iScript cDNA synthesis Kit (Bio-Rad) as the manufacturer’s protocol. The quantitative RT-PCR was performed using QuantStudio 7 Flex machine with Taqman probes for *TEAD1* (Hs00173359_m1), *TEAD2* (Hs01055894_m1), *TEAD3* (Hs00243231_m1), *TEAD4* (Hs01125032_m1), *RNF146* (Hs00258475_s1), CTGF (Hs00170014_m1) and *GAPDH* (Hs02786624_g1) (Applied Biosystems). Relative expression of each gene to *GAPDH* of target genes was assessed for at least 3 biological replicates.

#### In situ PLA

In situ PLA was performed with Duo-link reagents (Sigma) following the instructions from the product manual. Primary antibodies used in PLA were anti-TEAD4 antibody (Abcam 58310); RNF146 (HPA027209, Sigma-Aldrich); YAP (14074, CST); DAPI (4083, CST). Images were acquired with a Leica TCS SP5 confocal microscopy equipped with a CCD camera.

#### KGG ubiquitination site detection by mass spectrometry analysis

PTMScan® analysis: Cells were harvested and lysed in 20 mM HEPES pH 8.0, containing 9 M urea, 1 mM sodium orthovanadate, 2.5mM sodium pyrophosphate, and 1mM β-glycerophosphate. Whole-cell lysates from 4 conditions, control & overexpressed TEAD4 -/+ MG132 treatment (35 mg/condition) were subjected to reduction with 5 mM dithiothreitol (DTT) at 37 °C for 1h followed by alkylation with 10 mM iodoacetamide (IAA) in the dark at RT for 15 minutes. The protein mixture was diluted with HEPES buffer such that the final concentration of urea was 2M. Tryptic digestion was performed at an enzyme: substrate ratio of 1:50 at 37 °C overnight. After incubation, the digestion was quenched with 20% trifluoroacetic acid (TFA) and excipients were removed from the peptide mixture using C18 Sep-pak® column (Waters, Milford, MA). Peptides were eluted with 3 x 4 mL of 40% acetonitrile (ACN)/0.1% TFA. The eluent was lyophilized to complete dryness. Immuno-affinity isolation of peptides containing di-glycine remnant (-GG) on lysine residues was performed using antibody recognizing the KGG motif according to the PTMScan® protocol (Kim et al., 2011; Rush et al., 2005). PTMScan® Proteomics System was in-licensed from Cell Signaling Technologies (Danver, MA). http://www.cellsignal.com/services/peruse_licensing.html

Mass spectrometric analysis: Samples were reconstituted in solvent A (2% ACN/0.1% formic acid (FA)), and injected via an auto-sampler for separation by reverse phase chromatography on a NanoAcquity UPLC system (Waters, Dublin, CA). Peptides were loaded onto a Symmetry® C18 column (1.7 mm BEH-130, 0.1 x 100 mm, Waters, Dublin, CA) with a flow rate of 1 μL/minute and a gradient of 2% to 25% Solvent B (where Solvent A is 0.1% Formic acid (FA)/2% acetonitrile (ACN)/water and Solvent B is 0.1% FA/2% water/ACN) applied over 85 minutes, with a total analysis time of 120 minutes. Duplicate injections were made. Peptides were eluted directly into an Advance CaptiveSpray ionization source (Michrom BioResources/Bruker, Auburn, CA) with a spray voltage of 1.4 kV and were analyzed using a Q Exactive-HF mass spectrometer (ThermoFisher, San Jose, CA). Precursor ions were analyzed in the Orbitrap at 60,000 resolution; MS/MS data were acquired at 15,000 resolution. The instrument was operated in data-dependent mode whereby the top 10 most abundant ions were subjected for HCD fragmentation.

Bioinformatic data analysis: MS/MS spectra were searched using the search algorithm Mascot (Matrix Sciences, London, UK) against a concatenated target-decoy database (UniProtKBConcat 2016_06) comprised of the UniProt human protein sequences, known contaminants and the reversed versions of each sequence. A 50 ppm precursor ion mass tolerance and 0.05 Da fragment ion tolerance were selected with tryptic specificity, and up to 3 miscleavages were allowed. Variable modifications were permitted for methionine oxidation (+15.9949 Da), iodoacetamide adduct for cysteine residues (+57.0215 Da), and ubiquitin di-glycine remnant (-GG signature) on lysine residues (+114.0429 Da). Peptide assignments were first filtered to a 5% false discovery rate (FDR) at the peptide level followed by filtering at the protein level at 2 % FDR. The AScore algorithm was applied to a subset of peptides containing ubiquitinated species for site localization. Label-free quantitation of the KGG peptides across all four conditions was performed using XQuant, an algorithm that utilizes retention times to quantify peptides across runs (Bakalarski et al., 2008; Kirkpatrick et al., 2013). Linear mixed-effect modeling was performed for each protein to summarize peptide level changes. Significance values were calculated from the model, which considered replicate peptide measurements and repeat injections (Yu et al., 2015).

#### Project Achilles data analysis

Genome-wide CRISPR KO data (20Q1) was downloaded from depmap.org/portal/download. Univariate linear regressions were performed for all pairwise gene comparisons for 739 cancer cell lines. Gene-gene pairs with p-value < 10^-20^ were considered for visualization. Graph network included genes within 2 degrees of separation to *WWTR1* (TAZ) was visualized using igraph v. 1.2.5 R package. Communities were identified using random walks on a graph network.

#### Fly Lines and Husbandry

Flies were cultured on yeast/molasses-based food at 25°C or 29°C at 60% humidity with a 12-hour light/dark cycle. Female or male flies were used in all experiments. The following stocks were obtained from the Bloomington Stock Center (University of Indiana, Bloomington, IN): ey-gal4 (BL5535); luciferase TRiP RNAi (BL31603); y[d2] w[1118] P{ry[+t7.2]=ey-FLP.N}2; P{ry[+t7.2]=neoFRT}42D hpo[KC202]/CyO, P{w[+mC]=GAL4-Kr.C}DC3, P{w[+mC]=UAS-GFP.S65T}DC7 (BL25090); y[1] w[*]; P{ry[+t7.2]=neoFRT}42D P{y[+t7.7] ry[+t7.2]=Car20y}44B, P{w[+mC]=GMR-hid}SS2, l(2)CL-R[1]/CyO; P{w[+m*]=GAL4-ey.H}SS5, P{w[+mC]=UAS-FLP.D}JD2 (BL5251); P{ry[+t7.2]=neoFRT}42D; ry[605] (BL1802). The Hpo knockdown line was obtained from Vienna Drosophila Stock Center (Vienna Biocenter Core Facilities, Vienna, Austria) w1118; P{GD14028}v36028 (VDRC104169). The RNF146 overexpression line was obtained from FlyORF Zurich ORFeome Project (University of Zurich) M{UAS-RNF146.ORF.3xHA}ZH-86Fb (F003712). Recombinant lines ey-gal4 + UAS::hpoRNAi (BL5535 and VDRC104169) were generated using standard genetic methods.

#### Eye Imaging

Fly eye imaging was done using a LEICA M205FA stereoscope with the LEICA DFC 7000T camera. The light source used was a LEICA KL 1500 LCD fiber optic ring light. Flies were frozen at -80°C for two hours, heads were removed, and oriented using an apple agar plate.

#### Image and Statistical Analysis

The eye area (A.U.) was obtained using ImageJ by drawing an ROI around the eye. For sample size per genotype, see figure legend. One-way ANOVA with Sidak’s multiple comparison test was performed using PRISM (GraphPad PRISM 7). Because of the variance between both eye sizes of the same fly, we measured each eye individually. Statistical analysis was performed using data from all eyes of multiple individual animals.

#### TEAD Lipid Pocket TR-FRET Assay

His-tagged TEAD proteins were pre-incubated with TEAD project compounds for 30 minutes at room temperature. Biotinylated lipid pocket probes are then added to the TEAD/compound mixture and incubated for 60 minutes at room temperature. The lipid pocket probe competes with the test compound for the TEAD lipid pocket until equilibrium is reached. After 60 minutes, Europium labeled anti-His (Perkin Elmer # AD0110) and XL665 labeled streptavidin (CIS Bio 610SAXAC) are added to the TEAD/test compound/lipid pocket mixture and incubated for 30 minutes. TR-FRET values are then measured using an EnVision multi-label plate reader (Perkin Elmer Cat# 2104- 0010A.) If the lipid pocket probe binds to TEAD as expected, a TR-FRET signal results from the proximity of anti-His Eu and XL665. If a TEAD lipid pocket binder binds and displaces the lipid pocket probe, the disruption of the TEAD:probe interaction results in a decrease in the TR-FRET signal. The potency of compounds as TEAD lipid pocket binders is determined by the IC50 value generated using a non-linear 4-parameter curve fit.

#### Docking Calculations

Compound A was docked to PDE6δ using the Schrodinger Software Suite 2019-4 (Schrödinger, 2019). The protein structure (PDB ID: 5ML2) was prepared for docking using the protein preparation wizard in the Maestro program. Water molecules were removed from the structure, and missing sidechain atoms for residues B10 and B26 were added using the preprocessing module. The structure was optimized and minimized using default parameters. Docking was performed using the Glide program. The Glide docking grid was constructed in Maestro’s Receptor Grid Generation Module, and default parameters were used with the following two exceptions: (1) a van der Waals scaling factor of 0.8 was applied to the non-polar protein atoms to account for limited receptor flexibility and (2) a positional constraint was created to require docked ligands to have an atom located within 1 Å of the position of the chlorine atom in the 5ML2 structure ligand (the TEAD lipid pocket ligands often contain halogenated hydrophobic groups that are likely to bind in a similar position to the 5ML2 ligand chlorophenyl group). Poses from the docking calculations were analyzed for strain energy using the sdfMMConfAnalysis module of the Chemalot software package (Lee et al., 2017). The pose chosen from the docking study had the best combination of docking score and strain energy.

#### Cell viability assay

Cells were seeded at 500-1000/well on a 96-well plate for 24 hours. They were then treated with indicated siRNAs (final concentration of 25nM) or indicated inhibitors (with indicated concentration). Cell growth was assessed using CellTiter-Glo Luminescent Cell Viability Assay (Promega). All cell viability data was collected on an Envision luminescence microplate reader and calculated for at least 6 replicates per time point, per condition. IC_50_ for the inhibitors was determined by fitting the nonlinear regression curve generated by GraphPad Prism.

#### Crystal violet cell growth assays

Cells were seeded at 0.2 million cells per well in triplicate in 6 well plates for 24 hours. Cells were then treated with compounds at the indicated concentration and grown for 6 days. Following, cells were washed in PBS, fixed in methanol for 10 min, stained with 0.5% crystal violet for 30 minutes, washed with water, and air dried. Stained colonies comprising more than 30 cells were imaged using a scanner.

### *In vivo* mouse xenograft study

All mouse experiments were performed according to the animal use guidelines of Genentech, a Member of the Roche Group, conforming to California state legal and ethical practices. MDA-MB-231 shNTC or MDA-MB-231 shTEAD1-4, 10 million cells were implanted, with Matrigel, subcutaneously in the right flank region of female C.B-17 SCID.Bg (Charles River Laboratories) mice (n=35 each cell line; n=70 total). Mice were randomly assigned 10 mice per group when tumor volumes reached approximately 150 to 350 mm^3^. The mice were stratified into four groups (n=10/group) for treatment as follows: Group 1: MDA-MB-231 shNTC, 5% sucrose water; Group 2: MDA-MB-231 shNTC, 1 mg/ml Doxycycline; Group 3: MDA-MB-231 shTEAD1-4, 5% sucrose water; Group 4: MDA-MB-231 shTEAD1-4, 1 mg/ml Doxycycline. All water bottles were changed once (sucrose) or thrice (DOX) weekly. Tumor sizes and body weights were recorded twice weekly over the course of the study. Mice with tumor volumes ≥2000 mm^3^ or recorded body weight loss ≥20% were promptly euthanized. An MTD was defined as the dose in which <20% body weight loss or no animal deaths were observed. Tumor volumes were determined using digital calipers (Fred V. Fowler Company, Inc, Newton, MA) using the formula (*L* x *W* x *W*)/2. A generalized additive mixed model (GAMM) was then employed to analyze transformed tumor volumes over time (Liang, 2005; Lin and Zhang, 1999).

All mice were euthanized by 34 days post first dose, unless earlier needed due to mice reaching tumors, weight loss, or other humane endpoints. At the end of the study, tumors were collected after mice were euthanized for histopathological and *in vitro* analysis.

#### Protein Expression and Purification

TEAD2-YBD (A217-D447) was purified as described (Noland et al., 2016) with the exception that the proteins were treated with 2.5% hydroxylamine (Sigma) at pH 7.0 for one hour at room temperature prior to size exclusion chromatography. The final protein sample was concentrated to 5 mg/ml.

#### Crystallization

TEAD2-YBD crystals of the base-centered monoclinic space group *C*121 were grown at 19°C by hanging-drop vapor diffusion using a drop ratio of 1:1 protein:reservoir solution and streak seeding. The reservoir solution contained 200 mM Potassium/sodium tartrate pH 6.5 and 20% PEG 3350. To obtain co-crystal structures, crystals were soaked for five days in 2 μL reservoir solution containing a final concentration of 2 mM compound and 35% PEG 3350. Crystals were then flash-frozen directly in liquid nitrogen.

#### Data Collection and Structure Determination

X-ray diffraction data were collected at beamline 5.0.2 at the Advanced Light Source. Data were processed using autoPROC and elliptically truncated using STARANISO (Evans, 2006; Evans and Murshudov, 2013; Kabsch, 2010; Tickle, 2017; Vonrhein et al., 2011; Winn et al., 2011). All structures were solved by molecular replacement using Phaser (McCoy et al., 2007). There were two molecules in the asymmetric unit. Structures were rebuilt in Coot (Emsley and Cowtan, 2004) and subjected to iterative rounds of refinement and rebuilding using Phenix (Adams et al., 2010) and Coot. Data processing and refinement statistics are summarized in Supplemental Table 4.

#### RNA-seq and data analysis

Total RNA was extracted using the Qiagen RNAeasy kit with the on-column genomic DNA elimination. RNA was quantified with Qubit RNA HS Assay Kit (Thermo Fisher Scientific) and quality was assessed using RNA ScreenTape on 4200 TapeStation (Agilent Technologies). For sequencing library generation, the Truseq Stranded mRNA kit (Illumina) was used with an input of 100 nanograms of total RNA. Libraries were quantified with Qubit dsDNA HS Assay Kit (Thermo Fisher Scientific) and the average library size was determined using D1000 ScreenTape on 4200 TapeStation (Agilent Technologies). Libraries were pooled and sequenced on NovaSeq 6000 (Illumina) to generate 30 million single-end 50-base pair reads for each sample.

For RNA-seq data analysis, RNA-seq reads were first aligned to ribosomal RNA sequences to remove ribosomal reads. The remaining reads were aligned to the human reference genome (GRCh38) using GSNAP version “2013-10-10,” allowing a maximum of two mismatches per 75 base sequences (parameters: ‘-M 2 -n 10 -B 2 -i 1 -N 1 -w 200000 -E 1 --pairmax-rna=200000 --clip-overlap). Transcript annotation was based on the Ensembl genes database (release 77). To quantify gene expression levels, the number of reads mapped to the exons of each RefSeq gene was calculated.

#### Global proteome sample preparation

MDA-MB-231 cells were treated with DMSO and 0.5μM of Compound A/D/E for 0, 2 hours, 4 hours, 8 hours. Cell pellets were lysed in 8 M urea, 150 mM NaCl, 50 mM HEPES (pH 7.2), and complete-mini (EDTA free) protease inhibitor (Roche) by 15X passages through a 21g needle. Protein concentrations were then estimated by BCA assay (ThermoFisher Pierce, Rockford, IL). Disulfide bonds were reduced by incubation with 5 mM DTT (45 min, 37°C), followed by alkylation of cysteine residues by 15 mM IAA (30 min, RT Dark), and finally capped by the addition of 5 mM DTT (15 min, RT Dark). Proteins were then precipitated by chloroform/methanol precipitation and resuspended in a digestion buffer (8 M urea, 150 mM NaCl, 50 mM HEPES pH 8.5). Initial protein digestion was performed by additing of 1:100 LysC followed by incubation at 37°C for 3 hours. Samples were then diluted to 1.5 M urea with 50 mM HEPES (pH 8.5) before the addition of 1:50 Trypsin and incubation overnight at room temperature. Peptide mixtures were acidified and desalted via solid phase extraction (SPE; SepPak - Waters, Boston, MA).

Peptides were resuspended in 200 mM HEPES (pH 8.5) and a 100µg aliquot of peptides was mixed with tandem mass tags (TMT, ThermoFisher Pierce, Rockford, IL) at a label-to-protein ratio of 2:1. After 1 hour of labeling the reaction was quenched by the addition of 5% hydroxylamine and incubated at room temperature for 15 minutes. Labeled peptides were then mixed, acidified, and purified by SPE.

Labeled samples were separated by offline high pH reversed-phase fractionation using an ammonium formate-based buffer system delivered by an 1100 HPLC system (Agilent). Peptides were separated over a 2.1x150 mm, 3.5 µm 300Extend-C18 Zorbax column (Agilent) and separated over a 75-minute gradient from 5% ACN to 85% ACN into 96 fractions. The fractions were concatenated into 24 samples, of which 12 were analyzed for proteome quantification. Fractions were concatenated by mixing different parts of the gradient to produce samples that would be orthogonal to downstream low pH reversed-phase LC-MS/MS. Combined fractions were dried, desalted by SPE, and dried again.

#### Quantitative mass spectrometry and data analysis

nanoLC-MS/MS analysis was performed on an Orbitrap Fusion Lumos mass spectrometer (ThermoFisher, San Jose, CA) coupled to a Waters NanoAcquity UPLC (Waters, Milford, MA). Peptides were separated over a 100 µm X 250 mm PicoFrit column (New Objective) packed with 1.7 µm BEH-130 C18 (Waters, Milford, MA) at a flow rate of 500 nL/min for a total run time of 180 min. The gradient spanned from 2% Buffer B (0.1% FA/98%ACN/2% water) to 30% B over 155 minutes and then to 50% B at 160 minutes. For mass spectrometry analysis, peptides were surveyed within FTMS1 analyses (120,000 resolution, AGC = 1x106, maximum injection time [max IT] = 50 ms) and the top 10 peaks were selected for MS/MS ensuring that any given peak was only selected once in a 45 second window (ppm tolerance = 5 ppm). For dynamic exclusion, the “one precursor per charge state” was ON. For MS2 analysis, precursors were isolated using the quadrupole (0.5 Th window), fragmented using CAD (NCE = 35, AGC = 1.5x10^4^, max IT = 100 ms), and analyzed in the ion trap (scan speed = Turbo). Following MS2 analysis, the top 8 ions were simultaneously selected (synchronous precursor selection – SPS, AGC = 1.5x10^5^, max IT = 150 ms) and fragmented by HCD (NCE=55) before analysis in the Orbitrap (resolution = 50,000). Raw data files are available via the MASSIVE data repository using the identifier MSV000085331.

All mass spectrometry data was searched using Mascot against a concatenated target-decoy human database (downloaded June 2016) containing common contaminant sequences. For the database search a precursor mass tolerance of 50 ppm, fragment ion tolerance of 0.8 Da, and up to 2 missed cleavages. Carbamidomethyl cysteine (+57.0214) and TMT labeled N-terminus and lysine (+229.1629) were applied as static modifications, while methionine oxidation (+15.9949) was set as a dynamic modification. Peptide spectral matches for each run were filtered using line discriminant analysis (LDA) to a false discovery rate (FDR) of 2% and subsequently as an aggregate to a protein level FDR of 2%. TMT-MS3 quantification was performed using Mojave, with only those PSMs possessing isolation specificities greater than or equal to 0.7 considered for the final dataset. Intensities of each PSM were added to the peptide and then protein (proteome) level within R. Expression is reported as relative abundance, which is the measured intensity of any given channel divided by the total intensity for that protein.

#### ATAC-seq analysis

ATAC-seq reads were analyzed using the ENCODE ATAC-seq pipeline v1. Briefly, reads were trimmed of adapters by cutadapt (version 1.9.1, cite DOI:10.14806/ej.17.1.200) and mapped to hg38 by Bowtie2 (version 2.2.6, cite https://doi.org/10.1038/nmeth.1923). Bam files were converted to tagAlign format, which was then adjusted for Tn5 insertion sites by shifting +4bp for positive strand and -5bp for negative strand. TagAlign files were used to call peaks using MACS2 (version 2.1.0, cite DOI: 10.1186/gb-2008-9-9-r137), and those peaks with p<1e-6 were retained for differential analysis by Diffbind (version 3.0.13, cite http://www.nature.com/nature/journal/v481/n7381/full/nature10730.html ). Differential peaks were defined by absolute fold change > 1 and FDR < 0.01. The genomic distribution of differential peaks was annotated by ChIPpeakAnno (version 3.24.1, cite doi: 10.1186/1471-2105-11-237). Bigwigs corresponding to fold change against the background control were generated by MACS2 and used to generate heatmaps by the heatmap function of Deeptools (version 3.5.0, cite doi:10.1093/nar/gkw257). Motif analysis of differential peaks was performed using HOMER (version 4.10) with all peaks as the background. The peaks were assigned using Poly-Enrich (cite https://doi.org/10.1093/nargab/lqaa006) with distal enhancer-gene target links (>5kb from TSS) plus 5kb locus definitions. Gene set enrichment was searched against Gene Ontology, MsigDB Oncogenic, and Hallmark.

## SUPPLEMENTARY FIGURE LEGENDS

**Figure S1: (A)** Structure of TEAD4 DNA binding domain in complex with DNA (Shi et al., 2017). All predicted TEAD4 lysine ubiquitination sites identified from the LC-MS/MS in Figure 1 are labeled in magenta in the structures. **(B)** Structure of the TEAD4 YAP-binding domain (blue) in complex with YAP (green) (Mesrouze et al., 2017). All predicted lysine ubiquitination sites are labeled in magenta. **(C)** Geyser plots showing changes in ratio and significance of ubiquitination of the whole proteome under MG132 treatment versus DMSO. TEAD4 is highlighted by a red dot.

**Figure S2: (A, B)** Validations of the E3 ligase screen. A, a sub-list of candidate siRNAs (2 siRNA for each target) was identified from the siRNA libraries of Figure 2. The signal was normalized to the negative control, set at 100. B, Taqman RT-PCR assay for Hippo target genes of cells transfected with the indicated siRNAs. **(C)** TEAD4 ubiquitination is reduced upon knockdown of *RNF146*. To knock down *RNF146*, the HEK293 cell line was transfected with sg*RNF146* for 48 hours. TEAD4-Myc and V5-ubiquitin plasmids were then transfected for an additional 48 hours, and an *in vivo* ubiquitination assay was performed, as described in Figure 1. TEAD4 expression and ubiquitinated TEAD4 were detected with anti-Myc antibody. β-Actin served as a loading control (n=3). **(D, E)** OVCAR-8 cells were transfected with siNTC, si*RNF146* or si*YAP1* as indicated for 48 hours. RNA-seq was performed to measure *YAP1* and *RNF146* levels. **(F)** Taqman analysis showing upregulation of Hippo target gene expression upon *RNF146* knockdown in NCI- H226 cells and Detroit 562 cells. Cells were treated with siNTC (black bars), si*YAP1* (red bars) or si*RNF146* (green bars) for 72 hours before harvested for Taqman RT-qPCR. **(G-I)** *RNF146* knockdown enhanced TEAD4 half-life. G, OVCAR-8 cells were treated with siNTC or si*RNF146* for 48 hours. Note that 100 μg/mL of cycloheximide (CHX) or 10 µM of MG132 was added as a time course before cell harvest. The expression of indicated proteins was assessed by immunoblotting. β-Actin served as a loading control. H and I, H226, PA-TU-8902, Detroit 562, and OVCAR-8 cells were treated with si*RNF146* or sg*RNF146* for 48 hours. Note that 100 μg/mL of ChX was added for 15 hours before cell harvest. The expression of indicated proteins was assessed by immunoblotting. Levels of TEAD4 protein were compared at 15 hours after ChX was added. ** indicated t-test with p-value β 0.05 (n=3). Protein level quantification was done and normalized to siNTC (no ChX). β-Actin served as a loading control.

**Figure S3: (A, B)** RNF146 protein can be detected in the nucleus and cytoplasm of the cell. Detroit 562 cells were stained with RNF146 and NUP98 antibodies, as well as DAPI. Pan-TEAD and YAP were used as controls. (n=3). B, Detroit 562 and PA-TU-8902 were fractionated into nuclear and cytoplasmic compartments. Immunoblotting was performed with anti-RNF146, anti-YAP/TAZ anti-pan-TEAD, anti-Tubulin, and anti-b-Actin antibodies. **(C)** The schematic diagram for the PLA assay performed in Figure 2D using a TEAD4 antibody that detects both isoforms of TEAD4. Staining was observed in both nuclear and cytoplasmic compartments as expected. **(D)** RNF146 interacts with full-length TEAD4. PLA was performed using an anti-TEAD4 antibody (raised in mouse) and anti-RNF146 antibody (raised in rabbit) for detecting endogenous proteins in PA-TU-8902 cells, followed by imaging with confocal microscopy as described in the materials and methods section. Arrows indicate the foci of interaction (n=8). The PLA signals in Figure 2D were mostly found in the nucleus. **(E)** TEAD transcription factors interact with multiple members of the PARP family, including PARP1 and PARP9. IP mas-spec was done in PA-TU-8902 cell line using a pan-TEAD antibody which detects and pulls down all four TEAD paralogs. IgG antibody was used as a control for immunoprecipitation. The numbers represent peptide counts. Condition 1 was in the shNTC cell line, condition 2 was in the sh*YAP1/WWTR1* cell line; the interaction was detected in both conditions (n=1). **(F)** The ubiquitination of TEAD4 is significantly reduced upon the knockdown of *PARP1*. HEK293 cells were transfected with si*RNF146* or si*PARP1*. The cells were further transfected with TEAD4-Myc and V5-ubiquitin plasmids for 48 hours, followed by immunoprecipitation with an anti-V5 antibody. TEAD4 expression and ubiquitinated TEAD4 were detected with anti- Myc antibody (n=3). **(G)** D70A mutation enhanced TEAD4 half-life. MDA-MB-231 cells engineered to stably express sh*TEAD1/2/3/4* upon Doxycycline treatment were treated with Doxycycline for 24 hours to deplete endogenous TEADs, followed by transfection of TEAD4-Flag-Myc or TEAD4(D70A)-Flag-Myc and siNTC or si*RNF146* treatment for 48 hours. Note that 100 μg/mL of ChX was added as a time course before cell harvest. The expression of indicated proteins was assessed by immunoblotting. Levels of ectopic Flag-tagged TEAD4 and ectopic Flag-tagged TEAD4(D70A) proteins were compared and quantified before and after ChX was added (n=2). Protein level quantification was done and normalized to siNTC/TEAD4-Myc-Flag (no ChX). β-Actin served as a loading control.

**Figure S4: (A, B)** Knockdown of Scalloped results in a significantly smaller eye phenotype. A, the first group shows a control RNAi (ey:: Gal4>UAS:: luciferase RNAi) under an eye-specific driver (n=10). The second group shows the knockdown of Hpo (w1118; ey::Gal4/+: UAS::hpoRNAi/+) results in a significantly larger eye compared to the control. (n=10). The third group shows RNAi of Sd (ey:: Gal4>UAS:: SdRNAi) results in a significantly smaller eye phenotype (n=9). The fourth group show the RNAi of Sd in a hyperactive Yki background (hpoRNAi). It shows a similar phenotype of the RNAi of Sd (n=10). B, means +/- S.D. of eye area quantified per genotype are shown in arbitrary units (A.U.). One-way ANOVA with Sidak’s multiple comparisons. Numbers (n) of eye area are quantified and P-values are indicated above. **(C, D)** RNAi of RNF146 enhances the overgrowth phenotype caused by the hpo RNAi. C, the first group shows a control RNAi (ey:: Gal4>UAS:: luciferase RNAi) under an eye-specific driver (n=7). The second group shows knockdown of Hpo (w1118; ey::Gal4/+: UAS::hpoRNAi/+) results in a significantly larger eye compared to luciferase RNAi control. Also, the folding of the cuticle is seen behind the eye (n=10). The third group shows eyes RNF146 RNAi (ey:: Gal4>UAS:: RNF146RNAi) (n=10). The fourth group shows RNAi of RNF146 in a hyperactive Yki background (hpoRNAi) did not rescue the overgrowth phenotype and displayed an additive phenotype with a much larger eye (n=13). D Means +/- S.D. of eye area quantified per genotype are shown in arbitrary units (A.U.). One-way ANOVA with Sidak’s multiple comparisons. Numbers (n) of eye area quantified and P-values are indicated.

**Figure S5: (A)** Taqman RT-qPCR analysis showing downregulation of *TEAD1/2/3/4* and Hippo signature gene *CTGF* mRNA expression upon *TEAD1-4* knockdown in 6 Hippo pathway sensitive human cancer cell lines. Cells were treated with siNTC (black bars), and si*TEAD1/2/3/4* (red bars) for 72 hours before harvesting for Taqman. **(B)** Immunoblot analysis of NF2, pan-TEAD, YAP, TAZ, and β-actin upon *TEAD1/2/3/4* knockdown in Hippo pathway insensitive cell lines. **(C)** Body weight over time of mice bearing MDA-MB-231 cell xenograft tumors expressing *TEAD1/2/3/4* shRNA in the presence of sucrose (red) or doxycycline (green), or expressing NTC in the presence of sucrose (black) or doxycycline (blue). **(D)** Taqman analysis confirmed the knockdown of *TEAD1/2/3/4* in engineered MDA-MB-231 cells used for the xenograft study. **(E)** Chemical structure and biochemical data of TEAD-CIDEs screened in the study. **(F)** Abundance of different peptides identified with Compound D, Compound E, and Compound A as a function of time in proteomics analysis. Cells were treated with 0.5 μM of the compound for 0, 2 hours, 4 hours and 8 hours.

**Figure S6: (A)** Immunoblot of pan-TEAD, YAP, TAZ, and β-Actin as a function of time (0, 30 minutes, 2 hours, 4 hours, 8 hours, 24 hours) upon Compound D, Compound E, and Compound A treatment of MDA-MB-231 cells. **(B)** Immunoblot of pan-TEAD, YAP, TAZ, and β-Actin upon Compound B treatment of OVCAR-8 cells for 16 hours. **(C)** Docked structure of PDE68 with Compound A in the PDE68 structure (PDB 5ML2), Compound A in cyan, native PDB ligand (for comparison) in magenta. **(D, E)** PDE68 dependency from DepMap measured by Chronos score shows no correlation with Chronos scores of *WWTR1* and *NF2*.

**Figure S7: (A)** Cell viability data from 3D Cell Titer-Glo assay of Hippo pathway sensitive cell lines (in blue), black represents DMSO control-treated cells. **(B)** Cell viability data from 3D Cell Titer-Glo assay (in Relative luciferase unit) of Hippo pathway insensitive cell lines (in blue), black represents DMSO control-treated cells. The SK-N-FI growth response curve is shown in red.

## SUPPLEMENTARY TABLES

**Supplementary Table 1: KGG peptide counts for TEAD4**

**Supplementary Table 2: siRNA screen to identify negative regulators for the luciferase reporters driven by multimerized TEAD binding sites (8xGTIIC-Luc)**

**Supplementary Table 3: IP mass-spec in Patu-8902**

**Supplementary Table 4: X-ray crystallography data collection and refinement statistics**

