## Supplementary figures for "Targeting the Hippo pathway in cancers via ubiquitination dependent TEAD degradation"

**Figure S1**

A

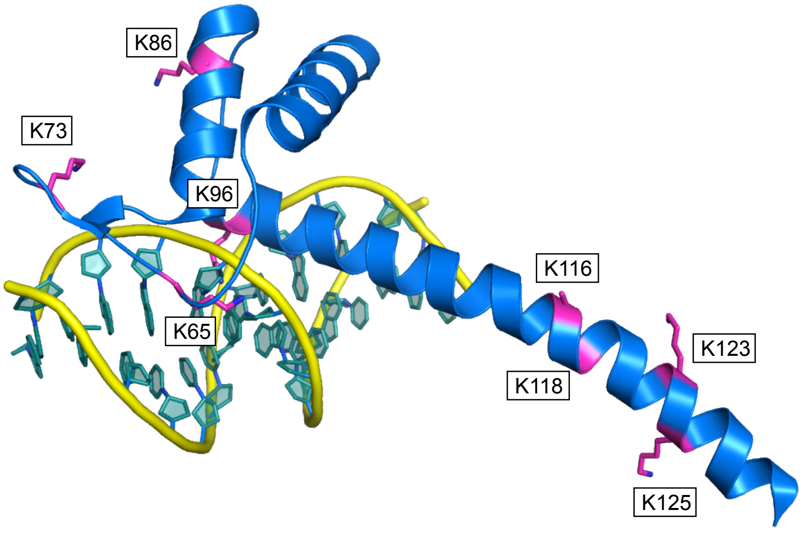


B

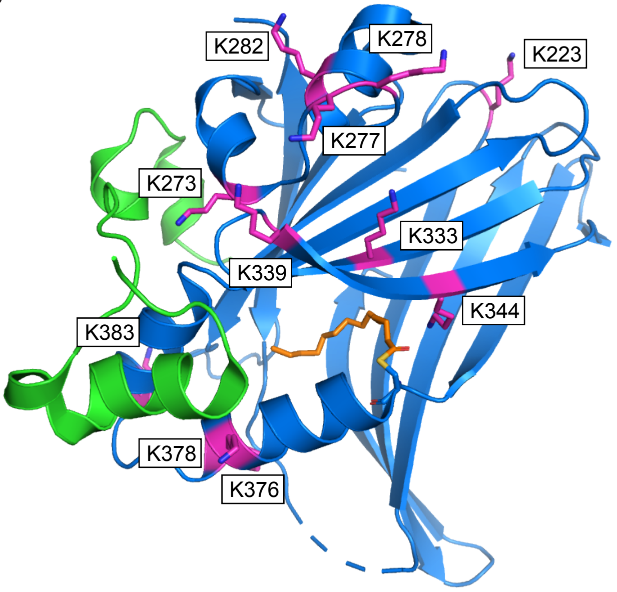


C

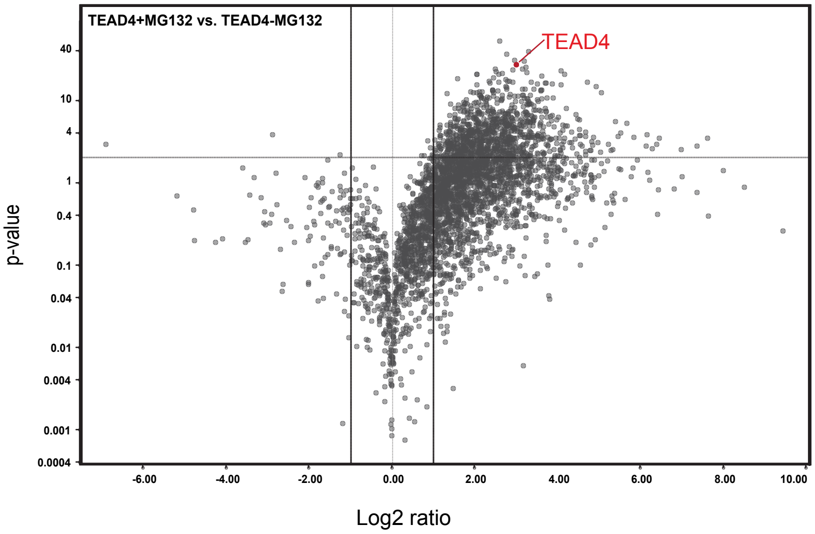

### Figure S2

**A**

| | | | | | | | | |
|---|---|---|---|---|---|---|---|---|
| siNeg#1 | siNeg#1 | siNeg#1 | UBL5-06 | UBL5-06 | UBL5-06 | RNF43-09 | RNF43-09 | RNF43-09 |
| siNeg#2 | siNeg#2 | siNeg#2 | UBL5-07 | UBL5-07 | UBL5-07 | RNF43-11 | RNF43-11 | RNF43-11 |
| RNF146-05 | RNF146-05 | RNF146-05 | WWTR1-s88 | WWTR1-s88 | WWTR1-s88 | YAP66 | YAP66 | YAP66 |
| RNF146-07 | RNF146-07 | RNF146-07 | WWTR1-s89 | WWTR1-s89 | WWTR1-s89 | YAP67 | YAP67 | YAP67 |
| TRAF3-11 | TRAF3-11 | TRAF3-11 | PHF5A-09 | PHF5A-09 | PHF5A-09 | PLK-1 | PLK-1 | PLK-1 |
| TRAF3-12 | TRAF3-12 | TRAF3-12 | PHF5A-11 | PHF5A-11 | PHF5A-11 | Q-Killer | Q-Killer | Q-Killer |
| 103.0 | 100.0 | 92.7 | 28.3 | 21.7 | 31.3 | 28.5 | 28.6 | 33.4 |
| 101.0 | 116.1 | 95.4 | 18.9 | 25.6 | 21.0 | 42.1 | 45.7 | 52.8 |
| 244.5 | 273.2 | 253.7 | 11.9 | 11.5 | 12.3 | 27.9 | 28.6 | 30.3 |
| 337.3 | 368.9 | 392.4 | 6.7 | 7.2 | 6.8 | 17.3 | 20.9 | 24.5 |
| 334.0 | 409.5 | 383.7 | 52.5 | 58.5 | 58.1 | 44.6 | 37.2 | 37.4 |
| 451.3 | 428.6 | 419.4 | 61.3 | 55.7 | 69.4 | 7.4 | 6.9 | 5.4 |

**B**

| RE, Gus, 72h | Gus | GapDH | CTGF | Cyr61 | BIRC5 | YAP1 | WWTR1 | KD |
|---|---|---|---|---|---|---|---|---|
| siNeg#2 | 1.00 | 1.00 | 1.00 | 1.00 | 1.00 | 1.00 | 1.00 | |
| NO TRMT | 1.00 | 0.81 | 1.54 | 0.98 | 0.86 | 0.66 | 0.94 | |
| siRNF43-09 | 1.00 | 0.56 | 0.54 | 0.67 | 0.49 | 0.51 | 0.60 | 0.00 |
| siRNF43-11 | 1.00 | 0.55 | 0.46 | 0.51 | 0.81 | 0.66 | 0.61 | 0.17 |
| siTRAF3-11 | 1.00 | 0.80 | 1.61 | 1.18 | 0.71 | 0.80 | 1.08 | 0.35 |
| siTRAF3-12 | 1.00 | 0.76 | 1.99 | 1.28 | 0.75 | 0.77 | 1.13 | 0.45 |
| siRNF146-05 | 1.00 | 0.72 | 2.58 | 1.33 | 0.72 | 0.61 | 0.96 | 0.16 |
| siRNF146-07 | 1.00 | 0.54 | 3.48 | 1.75 | 0.44 | 0.76 | 0.56 | 0.21 |
| siNeg#1 | 1.00 | 0.87 | 1.00 | 1.16 | 0.86 | 0.86 | 0.80 | |
| siYAP1 | 1.00 | 0.72 | 0.27 | 0.45 | 0.81 | 0.09 | 0.90 | 0.09 |
| siPHF5A-09 | 1.00 | 0.79 | 2.08 | 1.38 | 0.79 | 1.48 | 1.17 | 0.20 |
| siPHF5A-11 | 1.00 | 0.81 | 2.08 | 1.18 | 0.87 | 1.25 | 1.26 | 0.18 |
| siUBL5-06 | 1.00 | 0.85 | 1.34 | 1.18 | 0.68 | 1.14 | 1.06 | |
| siUBL5-07 | 1.00 | 0.76 | 1.22 | 1.00 | 0.63 | 0.95 | 1.16 | |
| siWWTR1-s88 | 1.00 | 0.47 | 0.42 | 0.46 | 0.54 | 0.72 | 0.11 | 0.11 |
| siWWTR1-s89 | 1.00 | 0.78 | 0.23 | 0.48 | 0.68 | 0.74 | 0.14 | 0.14 |

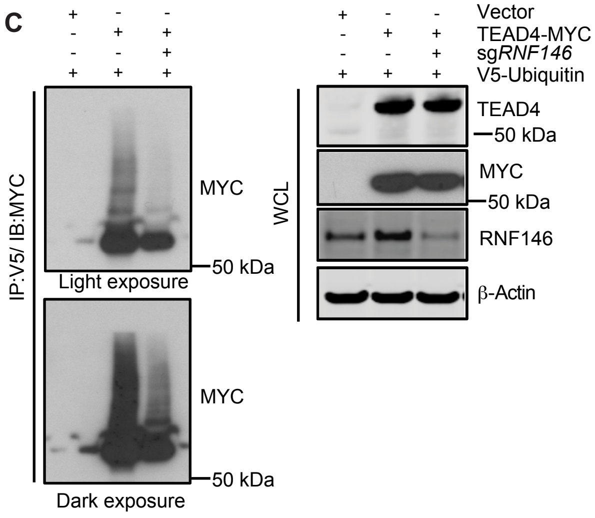


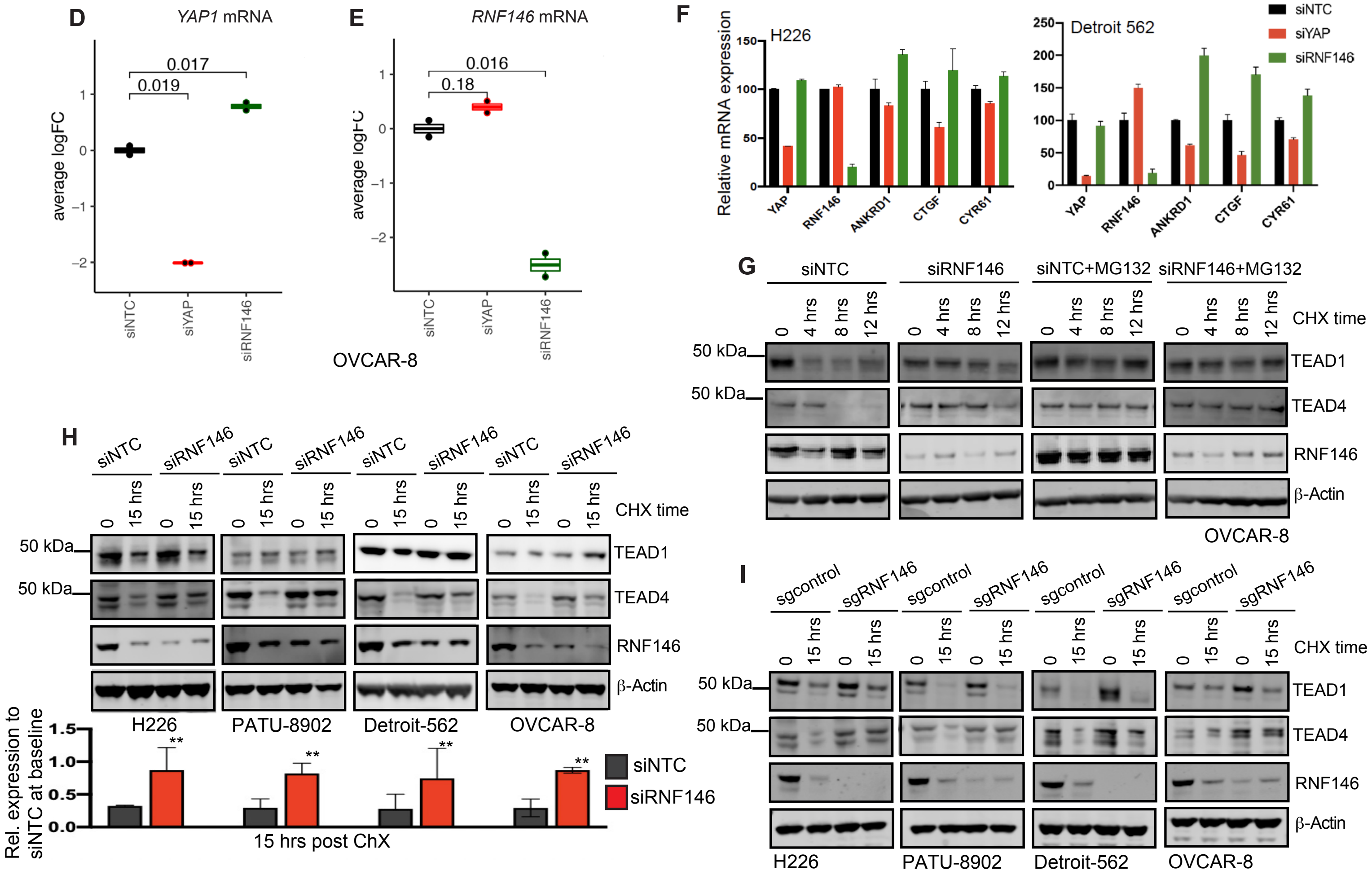

### Figure S3

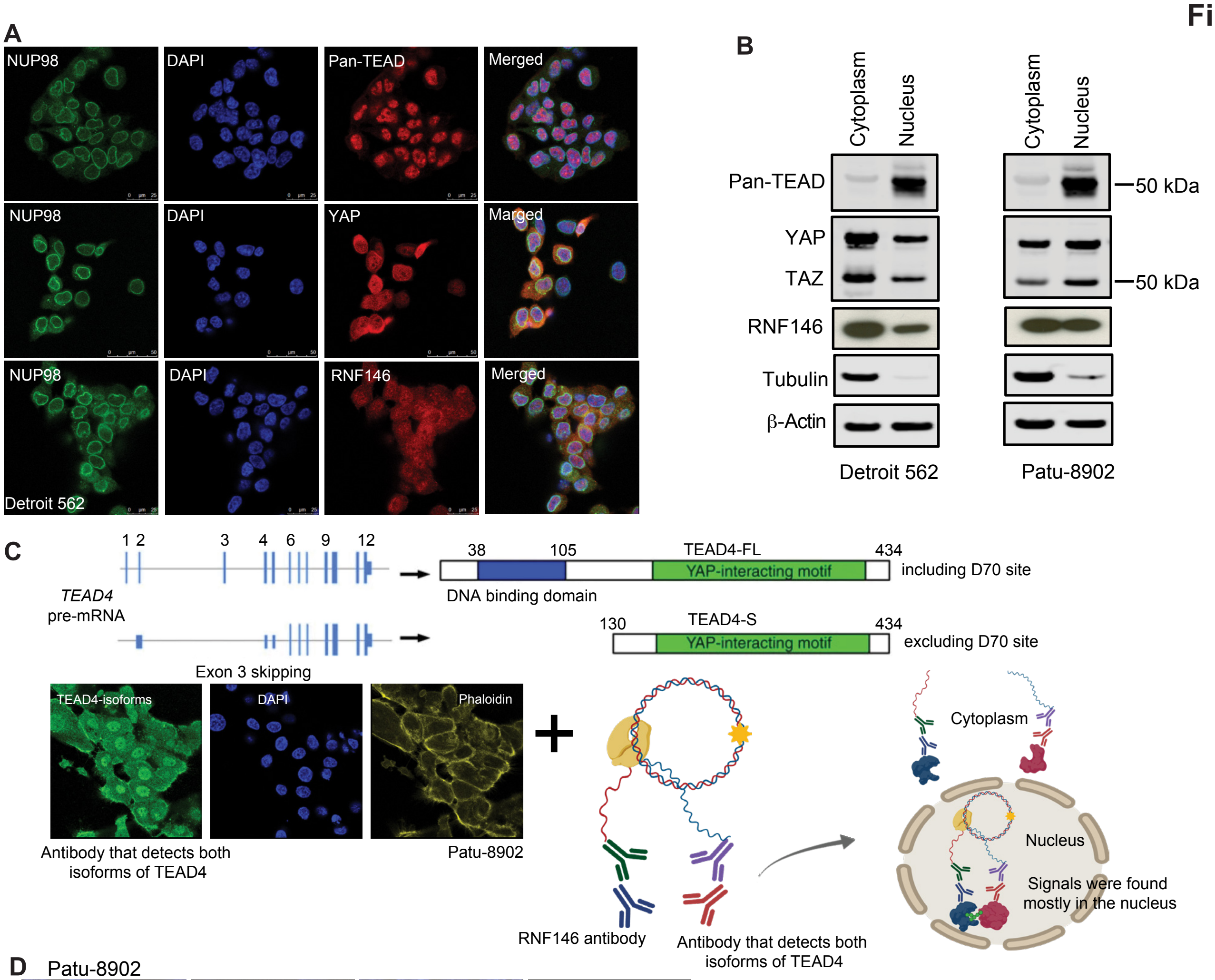


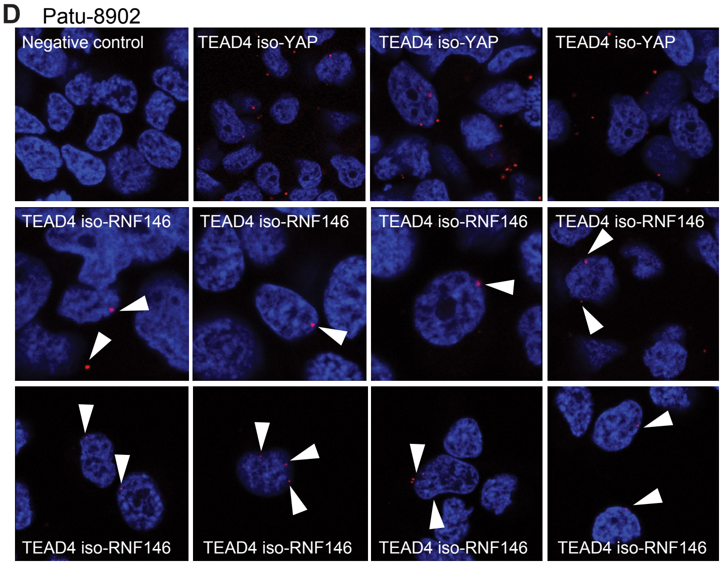


E

| Identifier | IgG_ condition1 | IP_ condition1 | IgG_ condition 2 | IP_ condition2 |
|---|---|---|---|---|
| PARP1_ P09874 | 0 | 52 | 0 | 26 |
| PARP9_ Q9UKK3 | 1 | 9 | 1 | 14 |

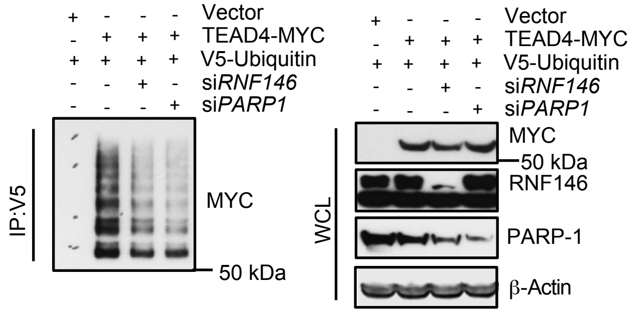


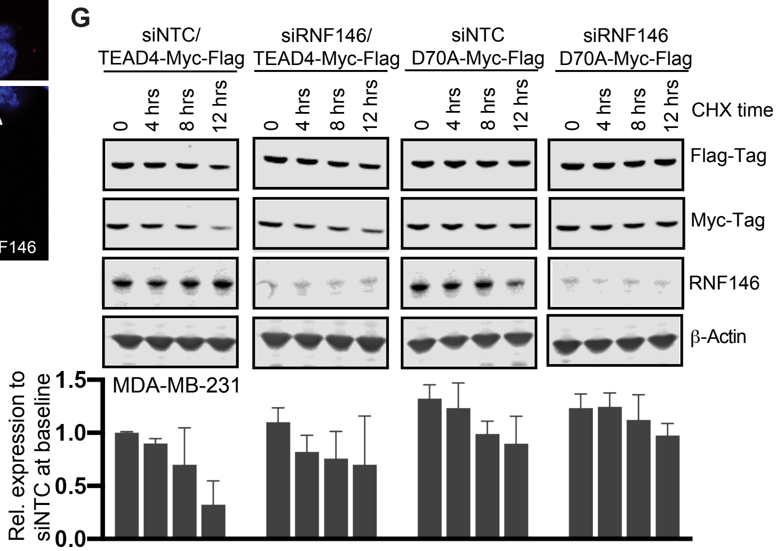

Figure S4

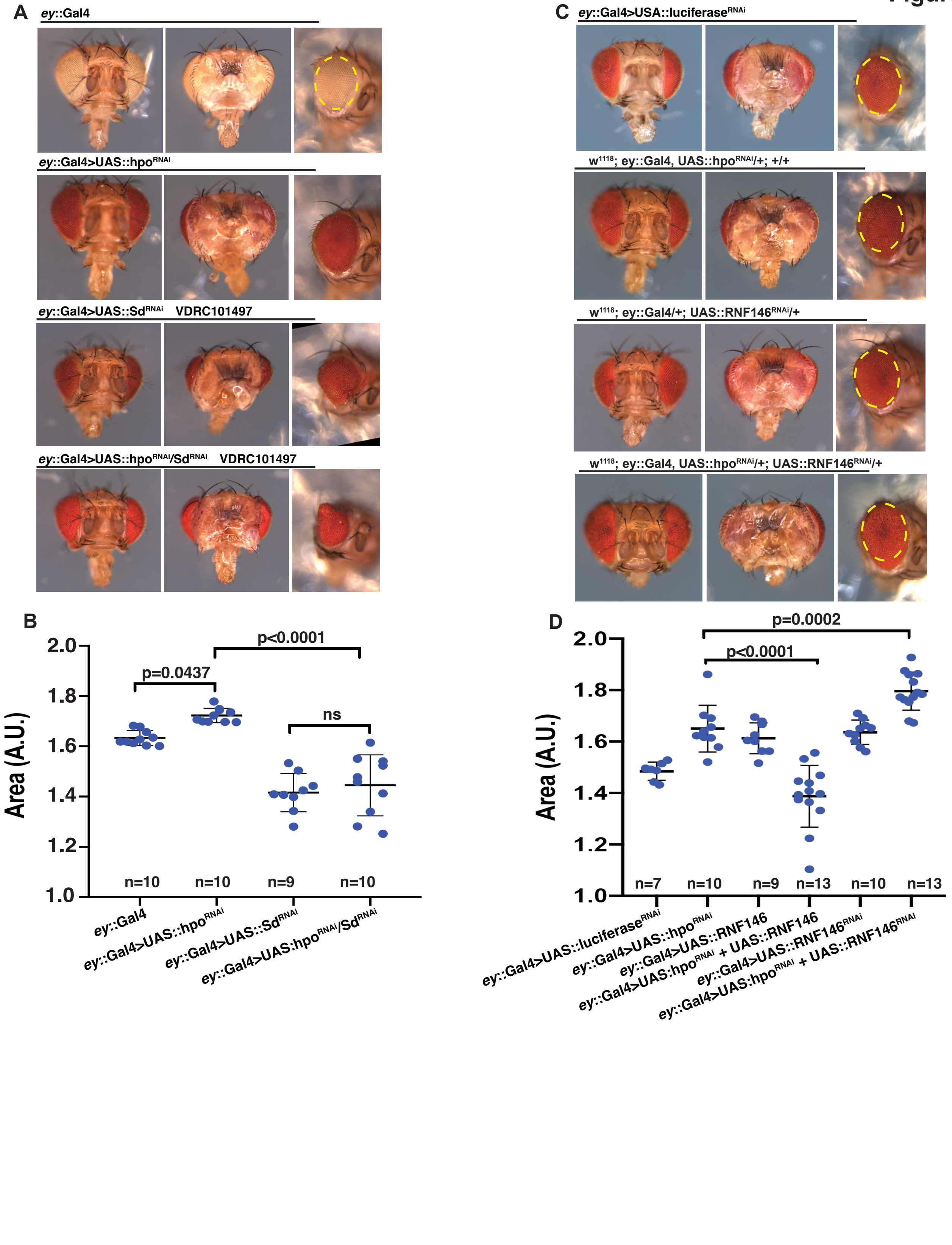

A
ey::Gal4
ey::Gal4>UAS::hpoRNAi
ey::Gal4>UAS::SdRNAi VDRC101497
ey::Gal4>UAS::hpoRNAi/SdRNAi VDRC101497
C
ey::Gal4>USA::luciferaseRNAi
w1118; ey::Gal4, UAS::hpoRNAi/+; +/+
w1118; ey::Gal4/+; UAS::RNF146RNAi/+
w1118; ey::Gal4, UAS::hpoRNAi/+; UAS::RNF146RNAi/+
B
Area (A.U.)
2.0
1.8
1.6
1.4
1.2
1.0
p=0.0437
p<0.0001
ns
n=10
n=10
n=9
n=10
ey::Gal4
ey::Gal4>UAS::hpoRNAi
ey::Gal4>UAS::SdRNAi
ey::Gal4>UAS:hpoRNAi/SdRNAi
D
Area (A.U.)
2.0
1.8
1.6
1.4
1.2
1.0
p=0.0002
p<0.0001
n=7
n=10
n=9
n=13
n=10
n=13
ey::Gal4>UAS::luciferaseRNAi
ey::Gal4>UAS::hpoRNAi
ey::Gal4>UAS::RNF146
ey::Gal4>UAS:hpoRNAi + UAS::RNF146
ey::Gal4>UAS::RNF146RNAi
ey::Gal4>UAS:hpoRNAi + UAS::RNF146RNAi

**Figure S5**

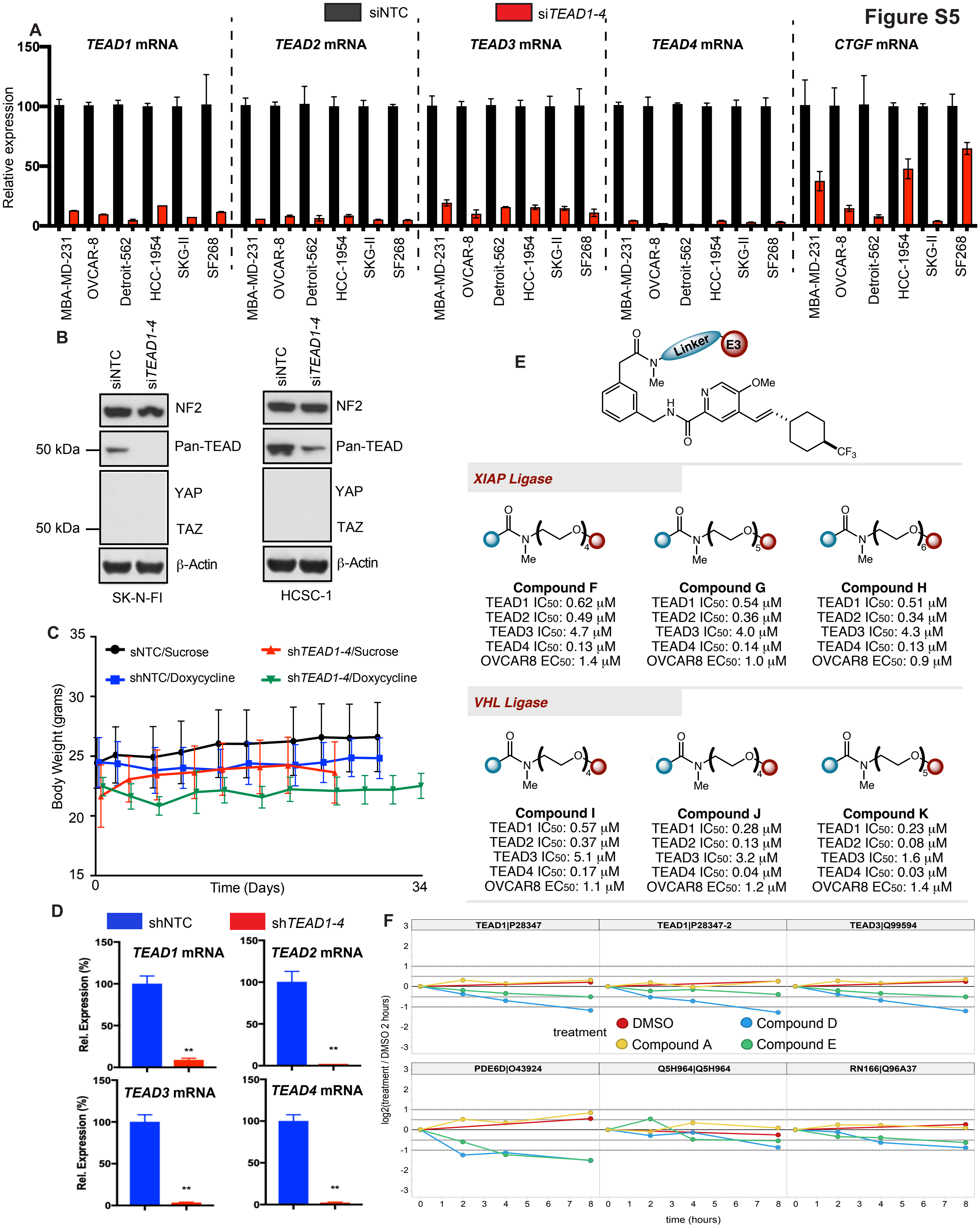

A
siNTC
siTEAD1-4
TEAD1 mRNA
TEAD2 mRNA
TEAD3 mRNA
TEAD4 mRNA
CTGF mRNA
Relative expression
150
100
50
0
MBA-MD-231
OVCAR-8
Detroit-562
HCC-1954
SKG-II
SF268
B
siNTC
siTEAD1-4
NF2
Pan-TEAD
YAP
TAZ
β-Actin
50 kDa
SK-N-FI
HCSC-1
C
sNTC/Sucrose
shTEAD1-4/Sucrose
shNTC/Doxycycline
shTEAD1-4/Doxycycline
Body Weight (grams)
Time (Days)
0
34
D
shNTC
shTEAD1-4
TEAD1 mRNA
TEAD2 mRNA
TEAD3 mRNA
TEAD4 mRNA
Rel. Expression (%)
E
Linker
E3
OMe
Me
CF3
XIAP Ligase
Compound F
TEAD1 IC50: 0.62 μM
TEAD2 IC50: 0.49 μM
TEAD3 IC50: 4.7 μM
TEAD4 IC50: 0.13 μM
OVCAR8 EC50: 1.4 μM
Compound G
TEAD1 IC50: 0.54 μM
TEAD2 IC50: 0.36 μM
TEAD3 IC50: 4.0 μM
TEAD4 IC50: 0.14 μM
OVCAR8 EC50: 1.0 μM
Compound H
TEAD1 IC50: 0.51 μM
TEAD2 IC50: 0.34 μM
TEAD3 IC50: 4.3 μM
TEAD4 IC50: 0.13 μM
OVCAR8 EC50: 0.9 μM
VHL Ligase
Compound I
TEAD1 IC50: 0.57 μM
TEAD2 IC50: 0.37 μM
TEAD3 IC50: 5.1 μM
TEAD4 IC50: 0.17 μM
OVCAR8 EC50: 1.1 μM
Compound J
TEAD1 IC50: 0.28 μM
TEAD2 IC50: 0.13 μM
TEAD3 IC50: 3.2 μM
TEAD4 IC50: 0.04 μM
OVCAR8 EC50: 1.2 μM
Compound K
TEAD1 IC50: 0.23 μM
TEAD2 IC50: 0.08 μM
TEAD3 IC50: 1.6 μM
TEAD4 IC50: 0.03 μM
OVCAR8 EC50: 1.4 μM
F
TEAD1|P28347
TEAD1|P28347-2
TEAD3|Q99594
PDE6D|O43924
Q5H964|Q5H964
RN166|Q96A37
treatment
DMSO
Compound D
Compound A
Compound E
log2(treatment / DMSO 2 hours)
time (hours)

#### Figure S6

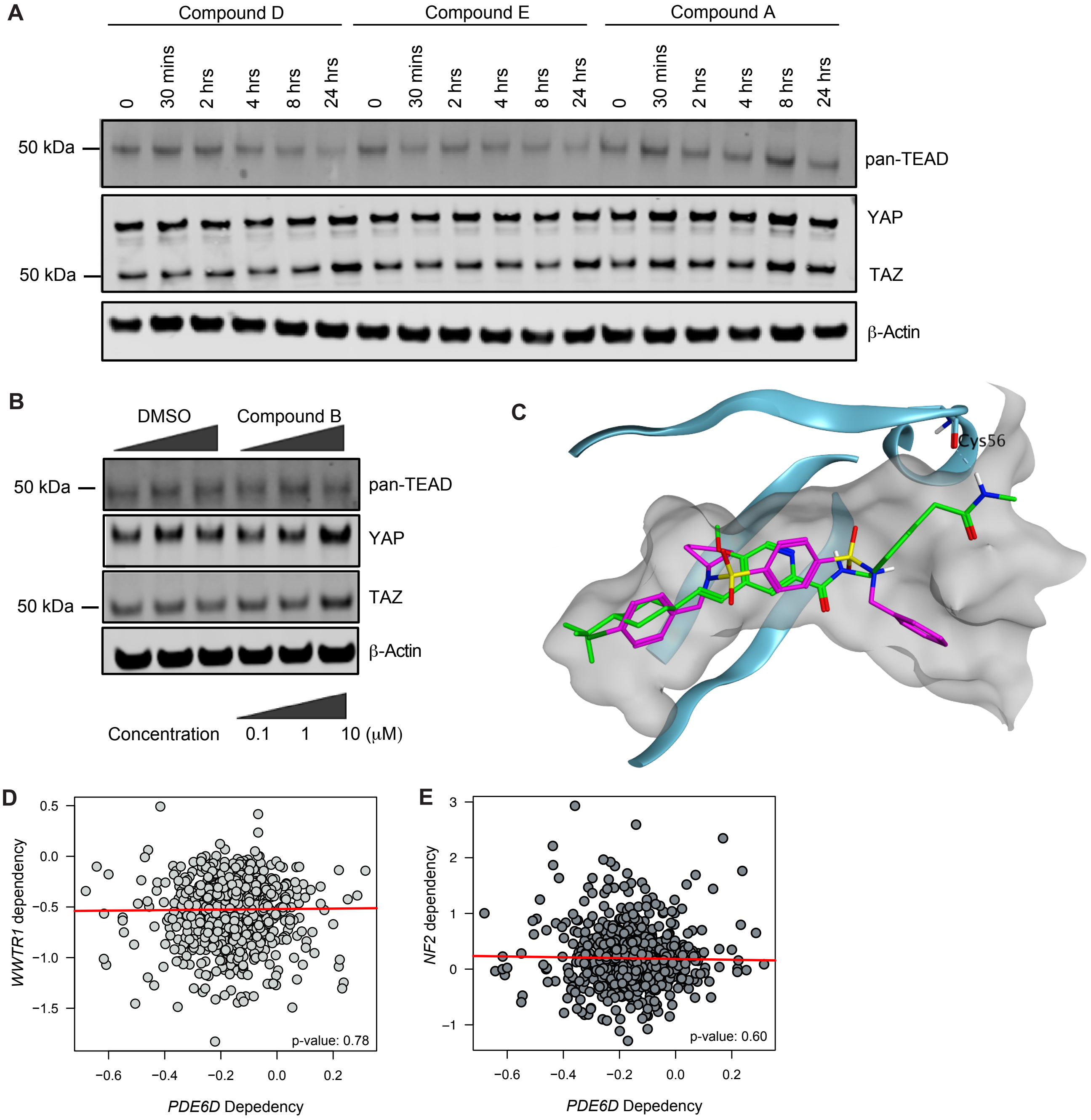

A
Compound D
Compound E
Compound A
0
30 mins
2 hrs
4 hrs
8 hrs
24 hrs
50 kDa
pan-TEAD
YAP
TAZ
β-Actin
B
DMSO
Compound B
50 kDa
pan-TEAD
YAP
TAZ
β-Actin
Concentration
0.1
1
10 (μM)
C
Cys56
D
WWTR1 dependency
PDE6D Depedency
p-value: 0.78
E
NF2 dependency
PDE6D Depedency
p-value: 0.60

### Figure S7

**A**

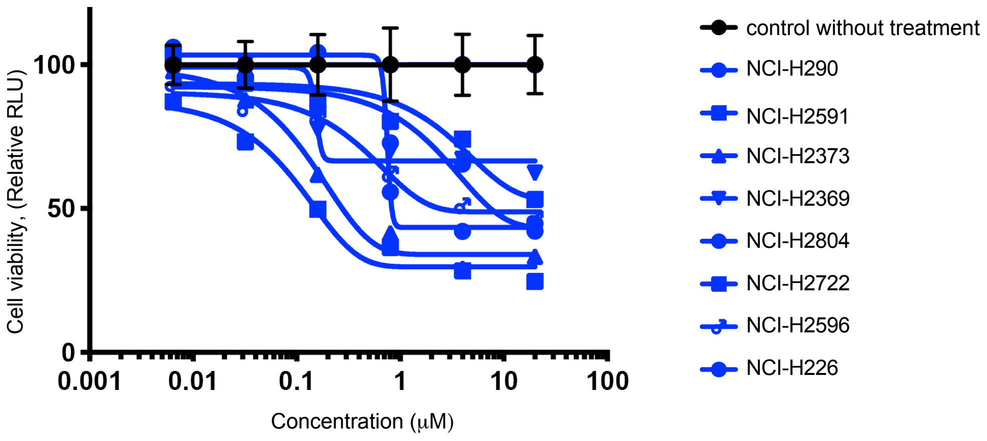

Cell viability, (Relative RLU)
100
50
0
0.001
0.01
0.1
1
10
100
Concentration (μM)
control without treatment
NCI-H290
NCI-H2591
NCI-H2373
NCI-H2369
NCI-H2804
NCI-H2722
NCI-H2596
NCI-H226


**B**

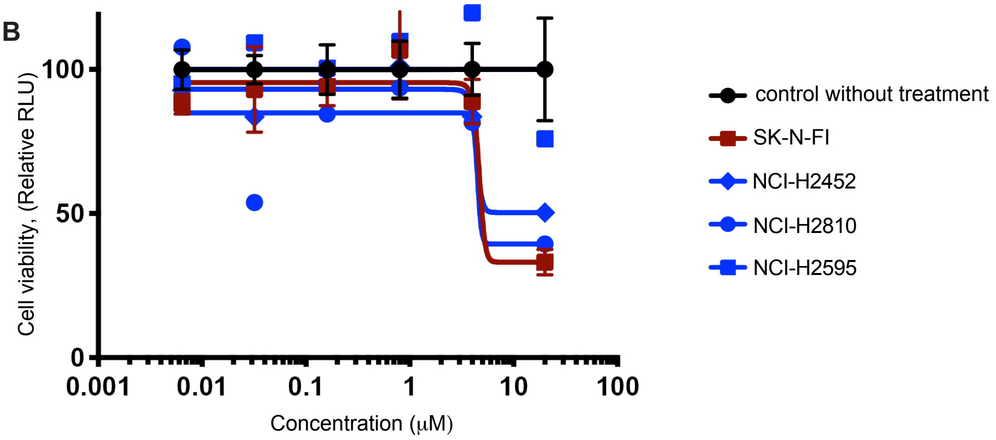

Cell viability, (Relative RLU)
100
50
0
0.001
0.01
0.1
1
10
100
Concentration (μM)
control without treatment
SK-N-FI
NCI-H2452
NCI-H2810
NCI-H2595
